# A spatial map of hepatic mitochondria uncovers functional heterogeneity shaped by nutrient-sensing signaling

**DOI:** 10.1101/2023.04.13.536717

**Authors:** Sun Woo Sophie Kang, Rory P. Cunningham, Colin B. Miller, Lauryn A. Brown, Constance M. Cultraro, Adam Harned, Kedar Narayan, Jonathan Hernandez, Lisa M. Jenkins, Alexei Lobanov, Maggie Cam, Natalie Porat-Shliom

## Abstract

In the liver, mitochondria are exposed to different concentrations of nutrients due to their spatial positioning across the periportal (PP) and pericentral (PC) axis. How these mitochondria sense and integrate these signals to respond and maintain homeostasis is not known. Here, we combined intravital microscopy, spatial proteomics, and functional assessment to investigate mitochondrial heterogeneity in the context of liver zonation. We found that PP and PC mitochondria are morphologically and functionally distinct; beta-oxidation was elevated in PP regions, while lipid synthesis was predominant in the PC mitochondria. In addition, comparative phosphoproteomics revealed spatially distinct patterns of mitochondrial composition and potential regulation via phosphorylation. Acute pharmacological modulation of nutrient sensing through AMPK and mTOR shifted mitochondrial phenotypes in the PP and PC regions, linking nutrient gradients across the lobule and mitochondrial heterogeneity. This study highlights the role of protein phosphorylation in mitochondrial structure, function, and overall homeostasis in hepatic metabolic zonation. These findings have important implications for liver physiology and disease.

## Introduction

Mitochondria are highly dynamic organelles that play critical roles in cell physiology, including energy production through oxidative phosphorylation (OXPHOS), metabolic signaling pathways, and biosynthesis ^1–5^. In response to changes in the environment, such as nutrient availability, mitochondria undergo remodeling through fission, fusion, and mitophagy ^2,4^. These dynamic rearrangements in mitochondrial architecture are fundamental to mitochondrial function, morphology and homeostasis ^4,5^. Further, these dynamic changes are modulated, in part, through post-translational modifications, like phosphorylation, allowing mitochondria to rapidly and reversibly adjust their metabolic output ^6–8^. *In vivo*, mitochondria in cells within organs are exposed to varying levels of nutrients due to their spatial positioning with respect to the blood supply. How this affects and/or regulates mitochondrial dynamics is not known.

This is particularly important in the liver, a central metabolic organ that balances whole-body nutrient availability. Within the liver, hepatocytes are organized into polygonal units called lobules. Nutrient-rich blood flows unidirectionally into the lobule, entering through the hepatic artery and portal vein (which will hence be referred to as periportal; PP) and drains into a single central vein (pericentral; PC). Consequently, the hepatocytes are exposed to different levels of regional metabolites and metabolic burdens depending on their relative location within the lobule ^9,10^. It has been previously shown that these gradients direct hepatocytes in different parts of the lobule to express different genes, a phenomenon known as liver zonation ^10^. Wnt ligands secreted by PC endothelial cells are a major driver of zonal gene expression ^11–13^. Although single-cell RNA sequencing has enhanced our understanding of liver zonation ^14,15^, the functional consequences are often inferred solely from gene expression.

Electron microscopy studies have revealed notable differences in mitochondrial morphology between PP and PC hepatocytes ^16–19^. These observations suggest that cells on the PP-PC axis, separated by up to a mere 300 μm, possess mitochondria with distinct functions. However, it is not known how spatial separation affects mitochondria functions *in vivo*. Furthermore, whether mitochondrial variations are determined genetically or continuously adjusted metabolically (via dynamic nutrient gradients) requires elucidation.

In the present study, we aimed to define the relationship between mitochondrial function, structure, and spatial positioning in the hepatic lobule. We also sought to identify mechanisms that regulate mitochondrial diversity during homeostasis. Our results highlight the role of protein phosphorylation and nutrient sensing in dynamically tuning zonated mitochondrial functions in the hepatic lobule. This is the first description of hepatic mitochondrial zonation, combining structural and functional features that reveal distinct subpopulations of mitochondria within the liver lobule.

## Results

### A comparative mitochondrial proteome of spatially sorted hepatocytes

To investigate how the spatial positioning of cells in the liver affects mitochondrial functions, PP and PC hepatocytes from the livers of four ad-lib-fed mice were enriched using unique surface markers and fluorescence-activated cell sorting (FACS; Fig 1A-C). E-cadherin and CD73 antibodies were used to enrich PP and PC hepatocytes, respectively, and Western blots were performed for validation (Fig 1D).

**Figure 1.**
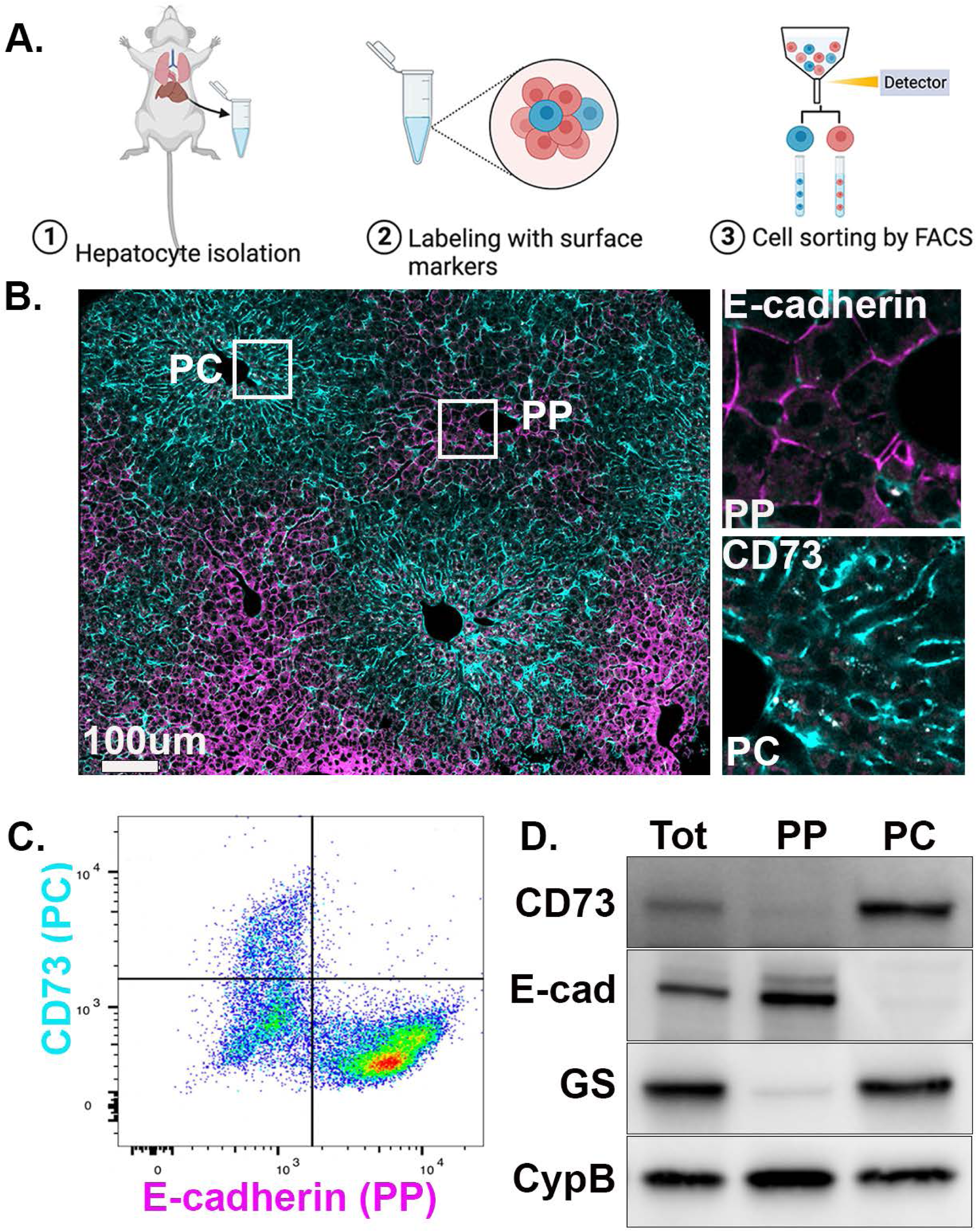
Spatial enrichment of PP and PC hepatocytes. **(A)** Schematic diagram depicting the workflow. Hepatocytes were isolated from the murine liver using two-step collagenase perfusion, after which unique surface markers were applied to label cells. Labeled cells were then enriched using fluorescence-activated cell sorting (FACS) to obtain hepatocytes from different zones. **(B)** Immunofluorescence staining of a liver section showing the zonal distribution of CD73 (cyan) and E-cadherin (magenta). **(C)** Two-dimensional scatter plot of hepatocytes labeled with CD73 and E-cadherin. **(D)** Western blot of spatially sorted hepatocytes. Abbreviations: Periportal (PP), pericentral (PC), glutamine synthetase (GS).

Next, PP and PC hepatocytes were subjected to tandem mass tag (TMT)-based quantitative mass spectrometry for total proteome analysis. The goal was to gain insight into mitochondrial functions by establishing a quantitative map of mitochondrial protein abundance along the PP-PC axis. Principal component and hierarchical clustering analyses showed sample grouping based on spatial origin (PP or PC) (Fig S1A-B). Of 5,018 proteins identified, 46% were zonated, meaning they had a biased expression toward PP or PC hepatocytes (Fig. S1C-D; Sup File 1). Pathway analysis highlighted PP and PC-restricted processes consistent with a previous study describing gene expression ^15^ (Fig S1E-F).

The list of quantified proteins was compared with the murine MitoCarta 3.0 database, which consists of 1,140 proteins. We identified 829 mitochondrial proteins, 422 of which were enriched in PP mitochondria and 113 in PC mitochondria (Fig 2A and B). To gain insight into the functions of PP and PC mitochondria, the top 25 unique proteins were selected, and their location and pathway within the mitochondria were determined using MitoCarta 3.0 database (Fig 2C and D). Selected mitochondrial proteins were also validated by immunofluorescence (Fig S2).

**Figure 2.**
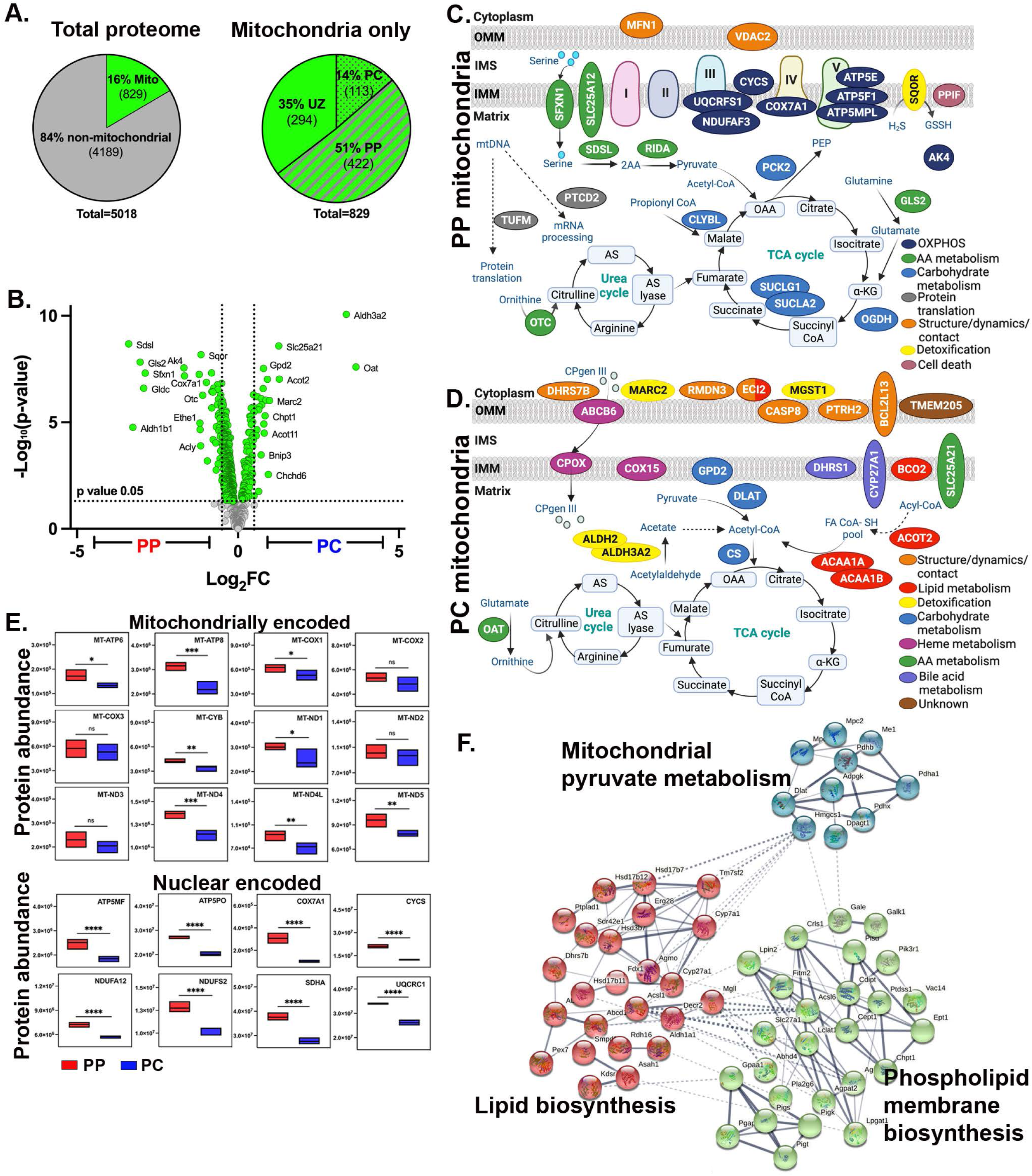
Comparative mitochondrial proteome of spatially sorted hepatocytes. **(A)** Total number of non-mitochondrial proteins (grey) and mitochondrial proteins (green) was detected by mass spectrometry (left). Percentage of periportal (PP), pericentral (PC), and unzonated (UZ) mitochondrial proteins (right); zonated expression based on a p-value (0.05). **(B)** Volcano plot shows the log_2_ PC/PP fold-change (x-axis) and the −log_10_ p-value (y-axis) for mitochondrial proteins. **(C-D)** Spatial distribution of the top 25 PP or PC mitochondrial proteins. Proteins were color-coded to reflect their cellular function. Pathways were listed based on the frequency at which they appeared. **(E)** Abundance of representative mitochondrial respiratory chain proteins, both nuclear and mitochondria encoded, are shown with box and whisker plots. **(F)** Bioinformatic STRING analysis of the PC mitochondria proteomic data. The interaction map illustrates the functional association of PC mitochondrial metabolism with cytosolic lipid synthesis. Data presented as mean⍰±⍰SD. *p < 0.05, **p < 0.01, ***p < 0.001, ****p < 0.0001.

Proteins enriched in PP mitochondria were primarily localized to the inner membrane or the mitochondrial matrix and were involved in amino acid metabolism or OXPHOS. Thus, we examined the spatial expression of selected OXPHOS proteins and found that PP mitochondria expressed higher levels of nuclear and mitochondrial-encoded OXPHOS components (Fig. 2E). In contrast, proteins enriched in PC mitochondria localized to the outer and inner mitochondrial membranes and matrix. In addition to regulating mitochondrial structure, dynamics, and contact with other organelles, many of the identified PC proteins are involved in lipid metabolism, detoxification, and carbohydrate metabolism (Fig. 2D). Notably, citrate synthase (CS), a tricarboxylic acid cycle (TCA cycle) enzyme, was highly expressed in PC mitochondria. In addition to the TCA cycle, when transported into the cytosol, citrate can be a precursor for lipid synthesis. Indeed, STRING analysis suggested a functional link between PC mitochondrial proteins involved in pyruvate metabolism and phospholipid membrane biosynthesis and lipid synthesis occurring in the cytosol (Fig 2F).

### Periportal mitochondria display enhanced bioenergetic capacity

We next performed a functional evaluation of the PP and PC mitochondria to determine their bioenergetic capacity. Intravital microscopy ^20^ was used to examine mitochondrial membrane potential in the intact liver of anesthetized mice. Mitochondria were labeled with MitoTracker green and TMRE, with the former labeling mitochondria matrix and the latter indicating membrane potential. TMRE labeled mitochondria in PP regions more intensely, indicating higher membrane potential (Fig 3A-B and Fig S3A-B). Isolated hepatocytes were labeled with Anti-CD73, Anti-E-cadherin antibodies, and JC1, a ratiometric fluorescent reporter of mitochondrial membrane potential. Consistent with the *in vivo* data, PP cells labeled in suspension had higher mitochondrial membrane potential than PC cells (Fig 3C and D). Together, these data show that PP mitochondria have a higher membrane potential that is not disrupted by spatial sorting, allowing the use of these cells for further physiological evaluation.

**Figure 3.**
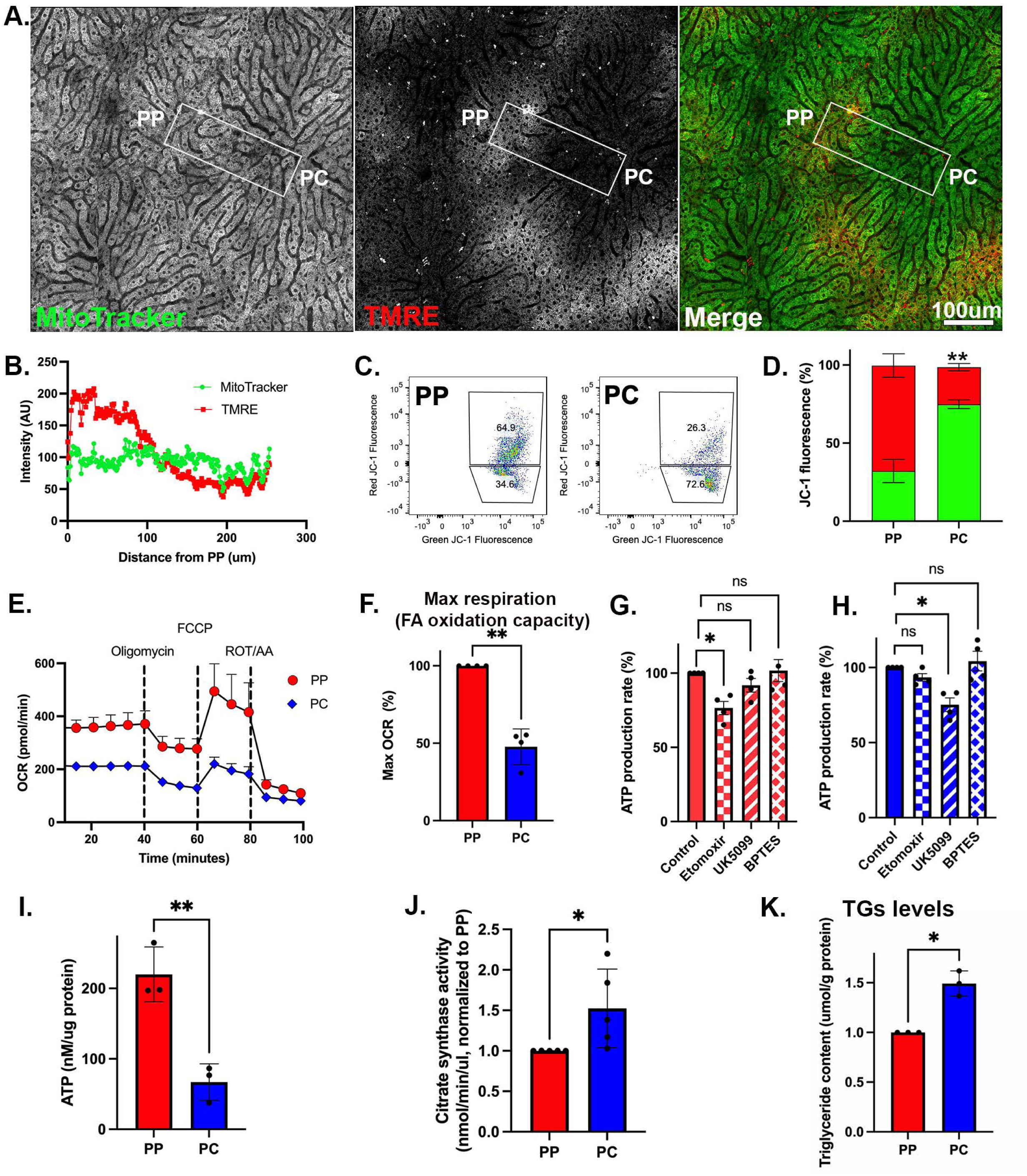
PP mitochondria display higher bioenergetic capacity. **(A)** Intravital microscopy of the hepatic lobule labeled with tetramethylrhodamine ethyl ester (TMRE), and MitoTracker Green to evaluate the relative mitochondrial membrane potential. The scale bar is 100 μm. **(B)** Mitochondrial membrane potential was evaluated by measuring fluorescence intensity (AU) along a PP-PC axis line. **(C-D)** Measurement of mitochondrial membrane potential using JC-1 and flow cytometry in spatially sorted hepatocytes. Dot plots of a representative experiment is shown. The bar graph shows three independent experiments. **(E)** Oxygen consumption rate (OCR) in spatially sorted hepatocytes using the XF Mito Stress Test Kit and Seahorse XF96 Analyzer. Samples were normalized to cell number. **(F)** Maximum respiration capacity in spatially sorted hepatocytes expressed relative to PP. The bar graph shows four independent experiments. **(G-H)** Substrate dependency assay in spatially sorted hepatocytes using the MitoFuel Flex test. ATP production rate relative to PP is shown in cells treated with etomoxir, UK5099, or BPTES. The bar graph shows the results of four independent experiments. **(I)** ATP content relative to PP in spatially sorted hepatocytes using the colorimetric luciferase assay from three independent experiments. **(J)** Citrate synthase activity relative to PP in spatially sorted hepatocytes from 5 independent experiments. **(K)** Intracellular triglyceride (TG) concentration relative to PP in spatially sorted hepatocytes from 3 independent experiments. Data presented as mean⍰±⍰SD; ns, *p < 0.05, **p < 0.01, ***p <0.001.

Subsequently, mitochondrial oxygen consumption rate and substrate preferences were evaluated using the Seahorse XF Analyzer in spatially sorted hepatocytes. PP hepatocytes consumed up to double the amount of oxygen compared to PC cells (Fig 3E) and had higher maximal respiration (Fig 3F). Next, substrate preference was determined by measuring the rate of ATP production in the presence of inhibitors. The capacity of PP mitochondria to produce ATP was significantly decreased by etomoxir, an irreversible inhibitor of fatty acid oxidation, whereas, in PC mitochondria, UK 5099, an inhibitor of the mitochondrial pyruvate carrier that inhibits pyruvate-dependent oxygen consumption, negatively impacted ATP production (Fig 3G and H).

ATP levels in PP hepatocytes were significantly higher as measured by a luminescent ATP Detection Kit (Fig 3I). On the other hand, citrate synthase expression and activity as well as triglyceride (TG) levels were higher in PC hepatocytes (Fig 2D and Fig 3J-K). Likewise, lipid droplets (LDs) which store TGs, were more abundant in PC regions of the lobule (Fig S4A). Higher TG levels in PC cells could be the result of lower lipid oxidation, increased lipid uptake, or decreased lipophagy. Alternatively, since high ATP levels inhibit citrate synthase activity ^21,22^, the lower ATP levels in PC hepatocytes may permit citrate synthase activity and lipogenesis.

Consistent with a previous report ^23^, mRNA and protein abundance of the lipogenesis-related enzymes Fasn (fatty acid synthase), Acly (ATP citrate synthase), Acaca (ACC1, acetyl-CoA carboxylase), and Scd1 (Stearoyl CoA desaturase 1) displayed a PP bias or were unzonated (Fig S4C-D). However, PP hepatocytes displayed higher levels of serine 79 phosphorylation, on acetyl-CoA carboxylase (ACC1), which inhibits lipogenesis supporting the hypothesis that lipogenesis preferentially occurs in PC hepatocytes (Fig S4B). This implies that the functional specialization of hepatocytes is not only regulated by differential expression but also via phosphorylation.

### Mitochondrial morphology and organization vary across the hepatic lobule

To better understand the spatial organization of mitochondria in the hepatic lobule, we used confocal microscopy to visualize mitochondria in liver sections from mito-Dendra2 mice ^24^. Consistent with previous reports ^16–19^, an apparent dichotomy in mitochondrial architecture was observed along the PP-PC axis, with short, round mitochondria in PP and tubulated mitochondria in PC hepatocytes (Fig 4A and Movie S1 and S2). The transition between these two phenotypes occurred in the mid-lobular area, with a limited number of cells containing both phenotypes within a single cell (Fig 4A).

**Figure 4.**
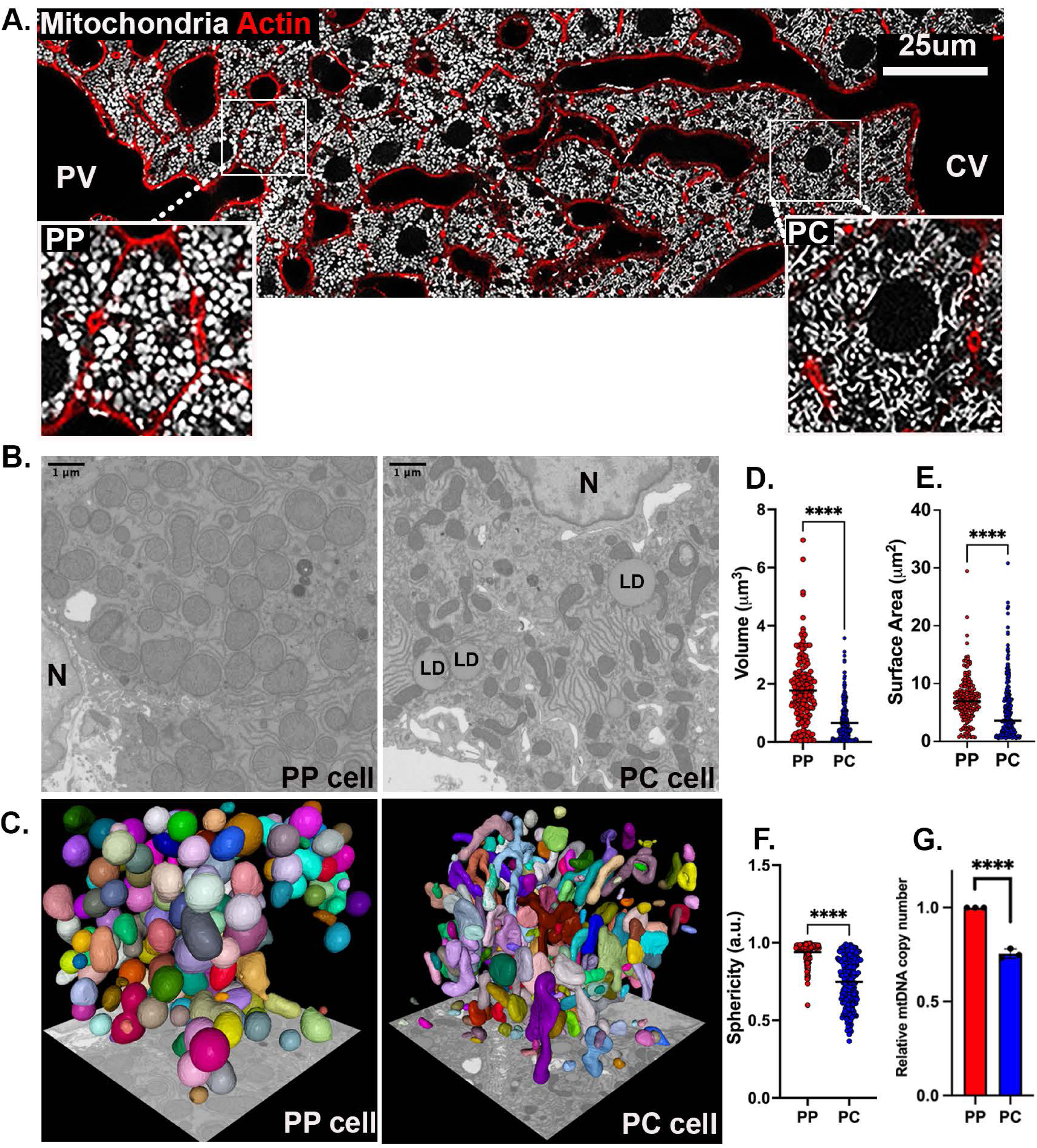
Mitochondrial morphology and organization across the lobule. **(A)** Confocal image of the PP-PC axis in liver sections from Mito-Dendra2 transgenic mice. Mitochondria are shown in white, and actin was labeled with phalloidin in red. Enlarged insets of a representative PP (left) and PC (right) cell are shown. **(B)** Mitochondria were visualized by Focused Ion Beam Scanning Electron Microscopy (FIB-SEM). Representative sections of PP (left) and PC (right) cells. Nucleus (N), Lipid droplet (LD). **(C)** Segmentation and volume rendering of mitochondria from PP (left) and PC (right) cells using Mito-Net and empanada-napari. **(D-F)** Quantification of mitochondrial morphological features including volume, surface area, and sphericity index (a measure of similarity to a perfect sphere (=1)). A collection of 175 mitochondria in PP and 250 in PC were analyzed. **(G)** Quantification of relative mtDNA copy number by qPCR in spatially sorted hepatocytes. The bar graph shows four independent experiments. Data presented as mean⍰±⍰SD. ** p < 0.01, ****p < 0.0001.

Mitochondrial topology was visualized in 3D and at nanometer resolution with Focused Ion Beam Scanning Electron Microscopy (FIB-SEM) ^25,26^. Initially, scanning electron microscopy images of the hepatic lobule were used to identify PP and PC regions for FIB-SEM (Fig S5). Next, FIB-SEM volumes of the selected cells were acquired, individual mitochondria segmented, and mitochondrial volumentric models generated (Fig 4B-C and Movie S3 and S4; see Material and Methods). Distinct subcellular organization of the mitochondrial network in different parts of the lobule were apparent (Fig 4B and C). Mitochondria volume and surface area in PP cells were approximately 2-fold greater than those measured in PC cells (Fig 4D and E). The sphericity index also significantly differed between PP and PC, with mean values of 0.93 and 0.74, respectively (Fig 4F). The higher mean values in PP hepatocytes indicate a shape closer to a sphere as demonstrated by the round surfaces rendered (Fig 4C).

Larger mitochondrial volume in PP hepatocytes, together with higher bioenergetic capacity (Fig 3) suggests that overall mitochondrial mass is higher in PP regions of the lobule. Lending further support to this idea, mitochondrial DNA copy number, a commonly used method to evaluate mitochondrial mass, was also higher in PP hepatocytes (Fig 4G). Mitochondrial structural diversity, including the typical PP and PC morphologies described above, was conserved in the human liver, suggesting a similar structure-function relationship may apply (Fig S6). Taken together, mitochondrial structural variations across the lobule strongly correlate with the functional diversity in lipid oxidation and biosynthesis under normal physiological conditions.

### Enhanced mitophagy flux in PC hepatocytes

Mitophagy, the selective degradation of mitochondria via autophagy, is stimulated in response to various signals, including hypoxia, nutrient deprivation, and glucagon signaling ^27,28^. In the liver, basal mitophagy is also activated by the daily feeding and fasting cycle ^29,30^. To evaluate if mitophagy is differentially regulated across the liver lobule, we applied intravital microscopy of mitochondria-targeted Keima in ad-lib-fed mice (mtKeima) ^31^. The mtKeima protein has different excitation wavelengths depending on the acidity of the mitochondrial environment; in a neutral environment, mtKeima excitation is at 440 nm, while in an acidic environment, excitation is at 560 nm (Fig 5A). The ratio of acidic-to-neutral excitation is a measure of mitophagy and the relative difference between mitophagy in the presence or absence of the protease inhibitor leupeptin, is used to determine mitophagy flux. Tile scans of entire lobules were acquired and ratios of acidic-to-neutral excitation were calculated in selected regions (Fig 5B). While in saline-treated mice, mitophagy was consistently higher in PP regions, leupeptin significantly increased mitophagy only in PC hepatocytes, suggesting a higher mitophagy flux in PC regions. (Fig 5C). Notably, the increase in mtKeima fluorescence in PP regions was driven by a higher signal in both the neutral and acidic compartments, while in PC regions it was mainly due to an increase in the acidic compartments (compare grey scale insets with and without leupeptin in Fig 5B). Similar trends were obtained with another mitophagy reporter Cox8-EGFP-mCherry ^32^ (Fig S7A and B). The spatial differences in mtKeima fluorescence intensities in PP and PC regions (Fig 5B) were not due to variations in mtKeima expression levels (Fig S7C).

**Figure 5.**
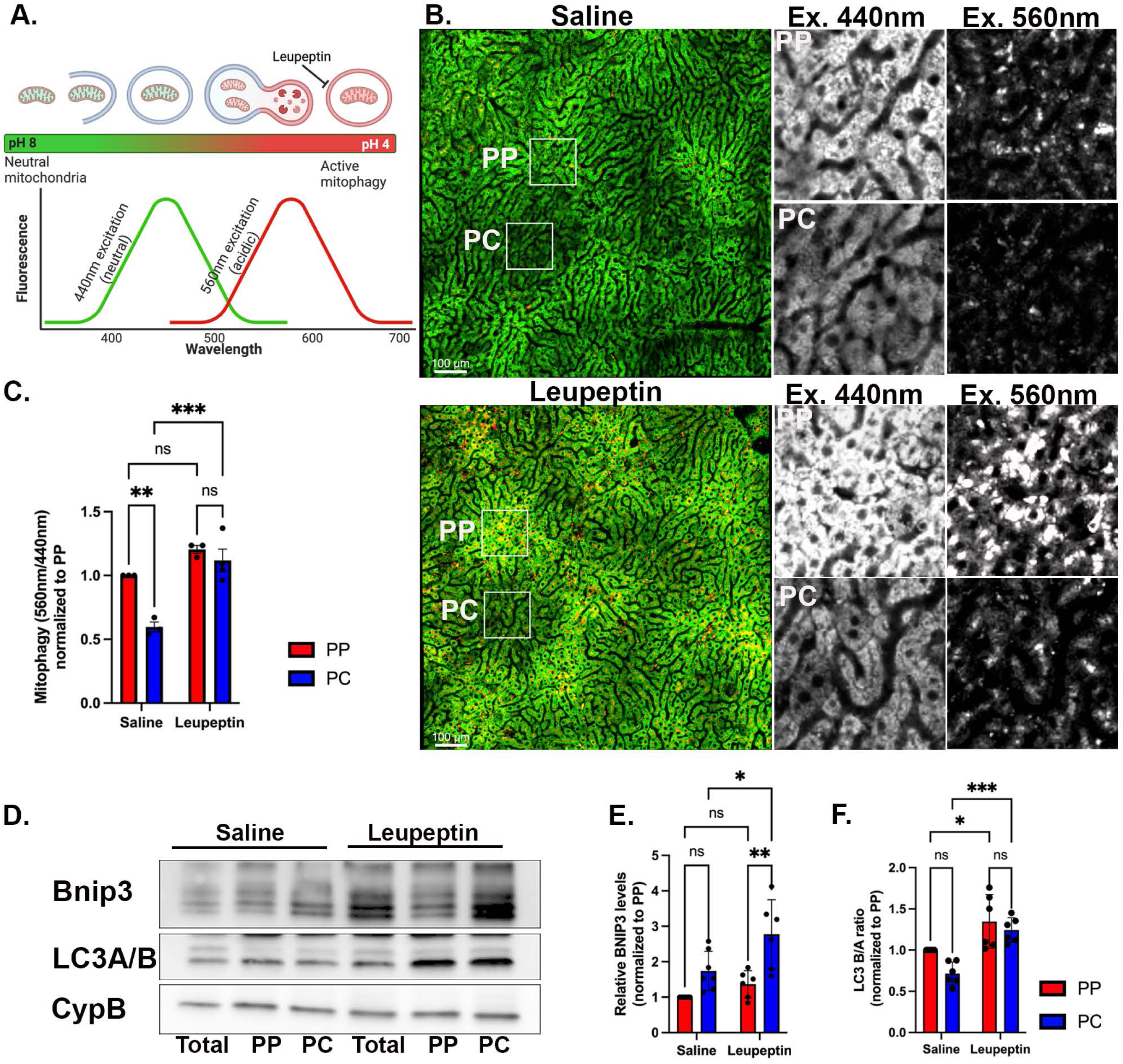
PC mitochondria display higher turnover via mitophagy. **(A)** mtKeima is a pH-sensitive mitophagy reporter. The ratio between the 440 nm (green; neutral pH) and 560 nm (red; acidic pH) excitation measures the proportion of mitochondria undergoing mitophagy. **(B)** Intravital microscopy of transgenic mice expressing mtKeima in saline-injected or leupeptin-injected mice. Representative images of the hepatic lobule and magnified insets of PP and PC regions **(C)** Quantification of mitophagy in PP and PC regions expressed as a fold change relative to PP cells. Bar graph shows four independent experiments. **(D)** Immunoblots of Bnip3 and LC3A/B of unsorted (total) and sorted PP and PC hepatocyte populations from livers treated with either saline or leupeptin. Mouse 1 (M1); Mouse 2 (M2) **(E-F)** Quantification of Bnip3 expression and LC3B/A ratio. Bar graphs show six independent experiments. Data presented as mean⍰±⍰SD. ns, *p < 0.05, **p < 0.01, ***p < 0.001.

We further tested the spatial differences in mitophagy by evaluating the levels of the mitophagy receptor Bnip3 and the autophagy marker LC3A/B using Western blots (Fig 5D). In saline-treated mice, there were higher levels of Bnip3 in PC hepatocytes, and significantly higher levels in mice treated with leupeptin, consistent with higher mitophagy flux in PC regions (Fig 5D and E). On the other hand, the proportion of LC3A/B (an autophagy marker) was similar in PP and PC hepatocytes, and leupeptin treatment had a comparable effect on both PP and PC cells, suggesting uniform levels of basal autophagy across the lobule (Fig 5DF). Other mitophagy-related proteins were enriched in PC hepatocytes (Fig S7D), as previously reported ^33,34^. Likewise, the lysosomal marker LAMP1 was distributed uniformly across the lobule, while LAMP1-positive lysosomes containing mitochondria were more abundant in PC hepatocytes (Fig S7E). Thus, basal mitophagy is higher in PC hepatocytes, which may be driven by the higher expression of mitophagy-related proteins and contribute to the lower mitochondrial mass.

### Phosphoproteome highlights the role of nutrient sensing signaling in shaping mitochondrial diversity

Reversible protein phosphorylation has been linked to cellular and mitochondrial metabolism ^6–8,35,36^. To examine the role of phosphorylation in regulating mitochondrial spatial heterogeneity and identify signaling pathways that may be involved, we profiled the phosphoproteomes of PP and PC hepatocytes (Fig S8A-C). We identified 7,278 phosphopeptides corresponding to 3686 phosphosites in 1,623 proteins (Fig 6A; Sup File 2). The majority of phosphorylation sites were on serine residues (89%), with a smaller proportion identified on threonine (10%) and tyrosine (1%) (Fig 6B). More than half of the proteins had a single phosphorylation site (Fig 6C). Interestingly, the overall number of phosphosites with PP or PC zonation was similar (Fig 6D), suggesting that ATP availability across the lobule (Fig 3A) is not limiting in homeostasis.

**Figure 6.**
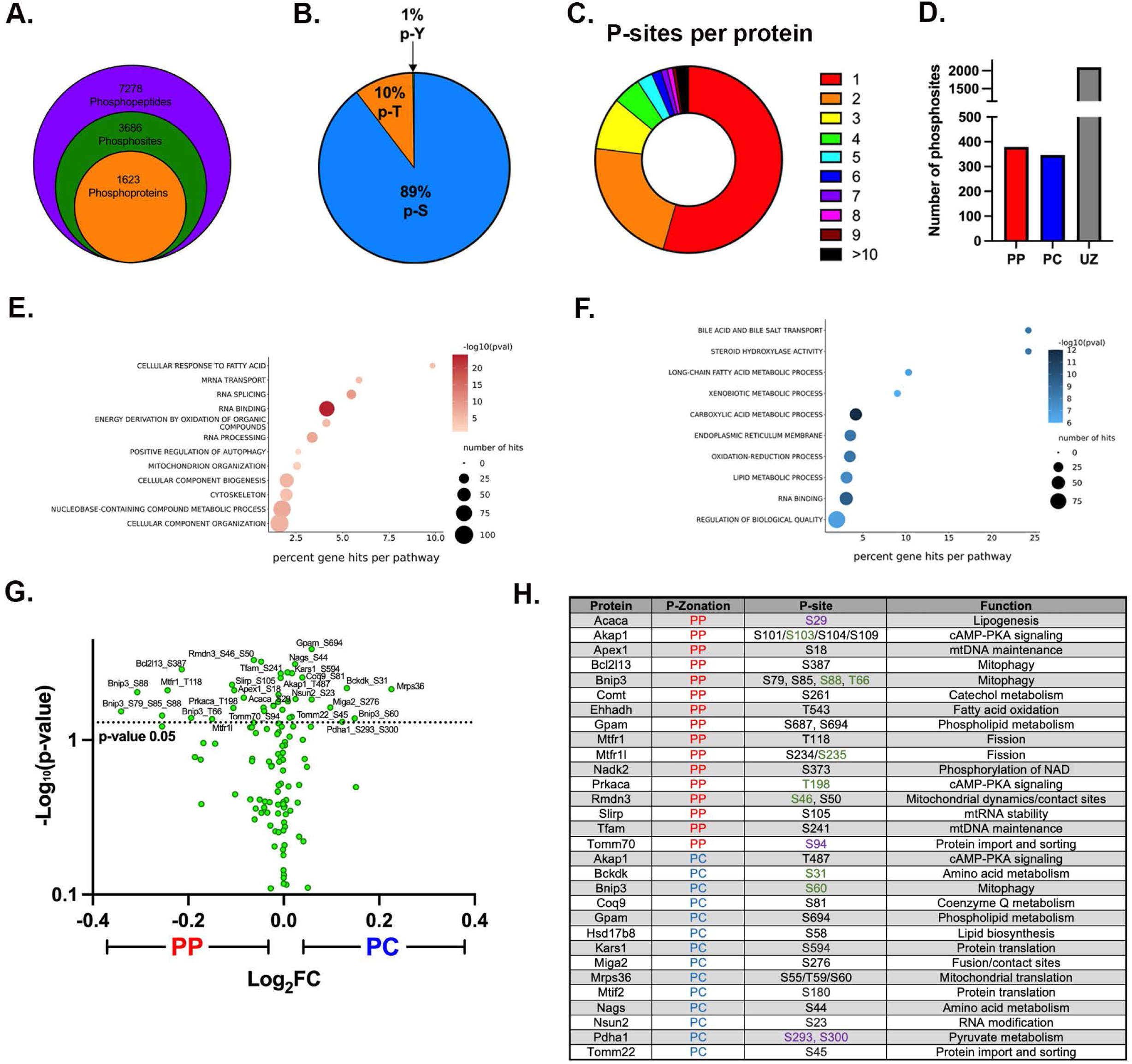
Phosphoproteome highlights spatially regulated pathways contributing to mitochondrial phenotypes. **(A)** Overview of phosphopeptides, phosphosites, and phosphoproteins identified in the phosphoproteome. **(B)** Proportion of phosphorylated serine (p-S), threonine (p-T), and tyrosine (p-Y) residues in the identified phosphopeptides. **(C)** Proportion of proteins identified with a certain number of phosphorylated residues per protein. **(D)** Number of phosphosites with a PP, PC, or unzonated (UZ) bias is shown in the bar graph. **(E-F)** The PP or PC zonated phosphoproteins were analyzed using GO enrichment analysis. **(G)** Mitochondrial phosphoproteome is shown in a volcano plot with log_2_ PC/PP fold-change (*x-axis*) and the −log_10_ p-value (y-axis). **(H)** Selected zonated mitochondrial phosphoproteins are shown in the table, including the phosphosites and the cellular function of the protein. Dash and comma marks distinguish between potential and identified sites, respectively. Activating phosphosites (green); Inhibitory (purple); uncharacterized (black).

Next, we performed a pathway analysis on zonated phospho-peptides to determine the processes that may be influenced by phosphorylation (either activating or inhibiting) in each hepatocyte population (Sup File 3). In PP hepatocytes, protein translation, metabolism, and positive regulation of autophagy were most notably regulated via phosphorylation (Fig 6E). On the other hand, multiple pathways related to lipid and phospholipid synthesis were represented in PC cells (Fig 6F). Sequence analysis of phosphorylated peptides using the online tool Momo (https://meme-suite.org/meme/tools/momo) revealed several phosphorylation motifs were enriched in different parts of the lobule, suggesting potential involvement of distinct kinases and signaling pathways (Sup File 4).

We also examined the phosphorylation of mitochondrial proteins, with mitochondrial phosphoproteome shown in the volcano plot (Fig 6G) and the zonated phosphoproteins for each hepatocyte group summarized in the table (Fig 6H and Sup File 5). Assessment of upstream regulators/kinases using PhosphoSitePlus highlighted factors such as leptin, insulin, mTOR, and AMPK as major drivers of the zonated phosphorylation (Sup File 6). To further examine the role of nutrient sensing in shaping the zonated phosphoproteome, we surveyed all zonated phosphopeptides for putative for AMPK and mTOR consensus sites using Group-based Prediction System (GPS) 5.0. Approximately 50% of the PP or PC phosphorylation sites were identified as putative AMPK or mTOR consensus sequences (Fig S8D). However, there was very little overlap in AMPK or mTOR substrates within PP and PC cells (Fig S8E). This suggests that although active throughout the lobule, AMPK and mTOR act on different substrates in PP and PC hepatocytes. To further substantiate this, we examined the phosphorylation status of two well-characterized mTOR substrates. Whereas phosphorylation on T389 of Ribosomal protein S6 kinase (S6K) displayed a PC bias, phosphorylation on T37/T46 of Eukaryotic translation initiation factor 4E (eIF4E)-binding protein 1 (4E-BP1), showed a PP bias, respectively (Fig S8F). Together, these results highlight a potential role for nutrient-sensing signaling in shaping zonated phosphoproteomes.

### Nutrient sensing signaling contributes to mitochondrial functional and structural heterogeneity across the lobule

Given the spatial variation in mitochondrial phospho-sites, their potential regulation by the nutritional state, and the capacity of mitochondria to respond to nutrient fluctuations ^1,8^, we hypothesized that nutrient-sensing signaling governs mitochondrial remodeling in hepatocytes. To test this, and decouple protein phosphorylation from transcription, we pharmacologically modulated AMPK or mTOR signaling *in vivo* and evaluated mitochondrial membrane potential (JC1), and lipid content (BODIPY) via flow cytometry (Fig 7B and C). Drug concentrations for acute response were determined by Western blotting of known downstream effectors (Fig S9A and B). Activation of AMPK (AICAR) and the inhibition of mTOR (Torin) were each associated with increased mitochondrial membrane potential in both PP and PC hepatocytes (Fig 7B). The inhibition of AMPK (Compound C; Cpc) and activation of mTOR (MHY1485) had no impact on membrane potential, suggesting the potential involvement of other factors (Fig 7B). On the other hand, the inhibition of AMPK, significantly increased lipid content in PC cells, and the activation of mTOR had a similar effect on PP cells, highlighting the spatial coordination of these pathways in lipid homeostasis (Fig 7C). Notably, the drugs had an acute effect on the mitochondria independent of cell size underscoring the complex spatial regulation of nutrient-sensitive signaling in the intact liver (Fig S9C).

**Figure 7.**
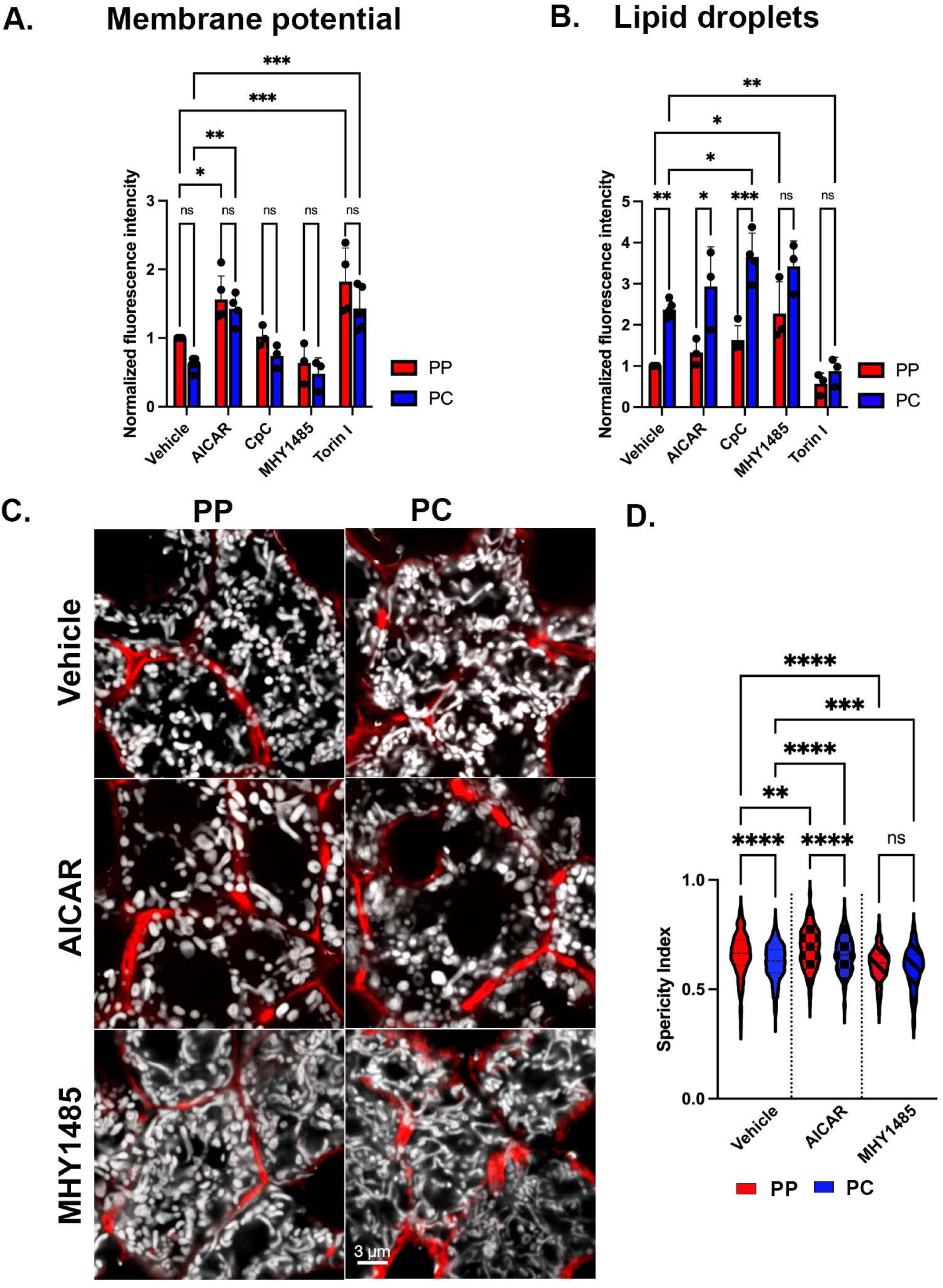
Nutrient sensing signaling contributes to mitochondrial functional and morphological diversity. **(A and B)** AMPK or mTOR signaling was modulated *in vivo* by injecting an activating drug, AICAR or MHY1485, or an inhibitory drug Cpc or Torin, respectively. The impact on mitochondrial membrane potential (JC-1) and lipid droplets (BODIPY) was evaluated using flow cytometry. Bar graphs show 3-5 independent experiments. **(C)** Confocal images of hepatocytes from liver sections of Mito-Dendra2 mice treated with vehicle, AICAR, or MHY1485. Mitochondria are shown in white, and phalloidin outlines hepatocytes in red. Scale bar= 3um. **(D)** Mitochondrial sphericity in mice treated with vehicle, AICAR, or MHY1485 was quantified. Data are presented as mean. ns, *p < 0.05, **p < 0.01, ***p < 0.001, ****p < 0.0001.

Finally, we performed microscopy studies in liver sections to examine if acute modulation of AMPK or mTOR impacted mitochondrial morphology. PC mitochondria from AICAR-treated mice had a round morphology with a higher sphericity index, similar to PP mitochondria in vehicle-treated cells (Fig 7C and D). Conversely, mTOR activation induced dense, elongated mitochondria in PP hepatocytes, with a lower sphericity index resembling the phenotypes observed in PC hepatocytes (Fig 7C and D). Taken together, the results show that mitochondrial zonation is acutely remodeled *in vivo* by nutrient-sensing signaling suggesting the nutrient gradient fine-tune mitochondrial metabolic output.

Mitochondrial heterogeneity can be established through various mechanisms, including differential gene expression (i.e. developmentally), dynamic gradients (i.e. metabolically), or a combination. Correlation between Wnt-regulated genes and the mitochondrial proteomes of both PP and PC were examined. We found a moderate negative correlation with the PP mitochondrial proteome and a moderate positive correlation with the PC mitochondrial proteome (Fig S9D). Overall, these experiments show that acute modulation of nutrient-sensing pathways shifts mitochondrial functions to impact mitochondrial functional diversity. These pathways work in concert with Wnt/β-catenin-regulated genes in determining the mitochondrial proteomes of both PP and PC.

## Discussion

In this study, we performed an in-depth characterization of mouse hepatic mitochondria along the PP-PC axis and assessed the spatial heterogeneity of these mitochondria in two populations of cells (Fig 1). We showed that mitochondria residing in hepatocytes adjacent to the portal vein produce significantly higher levels of ATP through lipid oxidation and OXPHOS. Roughly 300 μm apart, PC mitochondria oxidize pyruvate and produce higher citrate levels for lipogenesis. This PP-PC dichotomy in mitochondrial function mirrors the spatial separation of lipid utilization and biosynthesis in the liver. To our knowledge, this is the first spatial assessment of the molecular makeup and functional specialization of hepatic mitochondria. It is also the first in-depth investigation of how the physiological niche within the liver lobule alters mitochondrial metabolism. This location-dependent functional heterogeneity highlights the importance of considering mitochondrial disparity in understanding liver physiology and disease.

Consistent with previous observations ^16–19^, we also found striking variations in mitochondrial morphology across the PP-PC axis (Fig 4). Specifically, large spherical mitochondria correlated with enhanced OXPHOS in PP hepatocytes (Fig 2E and 4A-F). This morphology was previously shown to improve oxygen and nutrient exchange and tightly control calcium concentrations, an essential regulator of OXPHOS and mitophagy ^37,38^. Conversely, longer, tubulated mitochondria (Fig 4A-F) may protect against mitochondrial degradation ^37^, counterbalancing the higher mitophagy flux in PC regions (Fig 5A-C). Mitochondrial tubular versus spherical morphology may also impact mitochondrial metabolic tasks; with tubular mitochondria favoring enzymatic activities in the matrix, and spherical mitochondria improving bioenergetic efficiency. Shirihai and colleagues recently demonstrated that mitochondrial fragmentation, such as observed in PP hepatocytes, increases fatty acid oxidation ^39^. We further demonstrated a correlation between hepatic mitochondrial structure and function; pharmacological modulation of major nutrient signaling pathways resulted in a remodeled mitochondrial structure and its corresponding function (Fig 7).

Variations in mitochondrial fission and fusion are also likely to contribute to structural diversity, as has been previously shown in other tissues ^40,41^. We identified several candidates that may be involved in fusion pathways by regulating mitochondrial dynamics and contact sites in PC mitochondria (Fig 2D). On the other hand, mitochondrial morphology in PP hepatocytes suggests that fission is favored. Mitochondrial fission can be mediated by the phosphorylation of Mtfr1 or Mtfr1l, both of which showed a PP bias in our phosphoproteomic analysis (Fig 6H). Indeed, phosphorylation of the same conserved site in human Mtfr1l by AMPK was shown to cause mitochondrial fragmentation ^42^. Further experiments are necessary to examine the spatiotemporal regulation of mitochondrial fission-fusion dynamics Our finding that mitochondrial morphology is conserved in the human liver (Fig S6), suggests that some mechanisms are preserved across species and warrants further investigation.

Hepatic mitochondria not only display structural and functional heterogeneity, but they also possess distinct phosphoproteomes associated with each mitochondrial population. Despite higher ATP levels in PP hepatocytes (Fig 3I), the number of phosphosites was similar to in PC (Fig 6D). This suggests unique cytosolic or mitochondrial kinases or phosphatases are active in different cells. Indeed, we identified spatially distinct kinases reported to regulate the mitochondrial phosphoproteome ^6^, including Prakaca (PKA catalytic subunit) in PP mitochondria and, Pdha1 and Bckdk, in PC mitochondria (Fig 6H). Post-translational control of mitochondrial metabolism through phosphorylation has been shown to be critical across mouse strains, age, fasting/refeeding, and the onset of Type 2 diabetes ^35^. Unexpectedly, the inhibitory phosphorylation of Pdha1 was higher pericentrally (Fig 6H), which seemingly contradicts the Seahorse data showing that PC hepatocytes rely on pyruvate for ATP production (Fig 3H). However, the relative abundance of Pdha1 is also higher in PC hepatocytes (Sup File 1) which could lead to overall higher levels of the active enzyme. Our study adds to this complexity by demonstrating that differential regulation observed across the lobule is fundamental to hepatic cellular homeostasis. While these phosphorylation sites were not characterized in this study, combining the phosphoproteome data with functional studies allowed us to gain a system-level perspective into the functional consequences of protein phosphorylation (Fig 6).

Our research indicates that protein phosphorylation, in conjugation with zonated protein expression, plays a pivotal role in regulating various aspects of mitochondrial functionality, quality control, and lipid handling in distinct hepatic zones. The primary orchestrator of liver zonation at the transcriptional level is the Wnt/β-catenin signaling pathway ^12,13,43^. In agreement, we observed a correlation between the spatial distribution of mitochondrial proteins and genes regulated by the Wnt pathway (Fig S9D). However, we also found that mitochondrial proteins that display zonated phosphorylation, (Fig 6H), are intricately linked with spatially controlled pathways, including lipogenesis (Fig 3 and Fig S4), mitophagy (Fig 5 and Fig S7), and mitochondrial OXPHOS (Fig 3). Based on these observations, we hypothesize that the establishment of liver zonation is not solely dictated by gene expression; instead, it can be dynamically remodeled, through nutrient signaling pathways and protein phosphorylation.

The regulation of mitophagy and lipogenesis through reversible phosphorylation is ideal. It can be remodeled by nutrient-sensitive kinases (Fig 7), thus providing a quick mechanism for mitochondria to adjust their metabolic output. Several phosphorylation sites on the mitophagy receptor Bnip3 identified here are known to promote mitophagy ^44–46^. We also demonstrate functional zonation of lipogenesis through periportal inhibition of ACC1 (Fig S4B and 6H). Although additional investigation into the complex spatial coordination of AMPK and mTOR is required, our study shows that these kinases operate on distinct substrates in PP and PC cells (Fig S8D-E), which has a direct impact on mitochondrial phenotypes. This is especially critical in the liver, given its role in maintaining whole-body glucose homeostasis. We speculate that post-translational modifications provide the flexibility required for these vital metabolic adjustments, compared to the relatively sustained process of gene expression. Although further experiments to test this hypothesis are needed, we predict that protein phosphorylation may be the critical link between the observed discrepancies in function versus protein or gene expression. This finding has significant implications beyond the liver for studies where function is inferred from expression.

Advances in omics approaches and microscopy technologies have opened up new avenues for research and discoveries in hepatic cell biology. Although liver zonation was described nearly a century ago, we are only now starting to unravel how liver anatomy and the gradient of factors within it affect liver function at the tissue level. Our study, for the first time, demonstrates the delicate interplay between the function, shape, and space of hepatic mitochondria. We reveal that protein phosphorylation is an additional regulatory layer, offering spatial and temporal control of hepatic activities. Other post-translational modifications are also likely at play. In the current study, due to the sample size required for phosphoproteome and functional assays, we were limited to analyzing only two groups of hepatocytes. Therefore, significant technological improvements are required to allow for further and deeper exploration of liver zonation. Such advances are already underway with single-cell proteomics ^47^, allowing increased spatial resolution in murine models and human samples. Ultimately, understanding how the local physiological niche affects mitochondrial metabolism may be relevant for the therapeutic modulation of these pathways in liver-related pathologies.

## Contact information

Natalie Porat-Shliom, Center for Cancer Research, National Cancer Institute. Building 10, Room 12C207 MSC 1919, Bethesda, MD 20892. Phone: 240-760-6847.

## Acknowledgments

We thank Dr. Wen-Xing Ding for the Cox8-EGFP-mCherry-AAV8 construct. Drs. Torn Finkel and Nuo Sun for Mt-Keima mice. We thank Drs. Sudipto Das and Thorkell Andresson from the CCR proteomics Facility at the Frederick National Laboratory for Cancer Research and members of the NCI LGI Flow Cytometry Core. We thank Aayush Bhatawadekar and Abhishek Bhardwaj for volume renderings and morphological calculations based on vEM mitochondrial segmentation. Dr. Daniel Feliciano acquired images in Figures 4A and S2A. We thank Pranali Pathare Mangat of 3P Scientific Communications for providing scientific editing support. We are grateful for the critical comments made by members of the Porat-Shliom lab.

---

**Figure S1.**
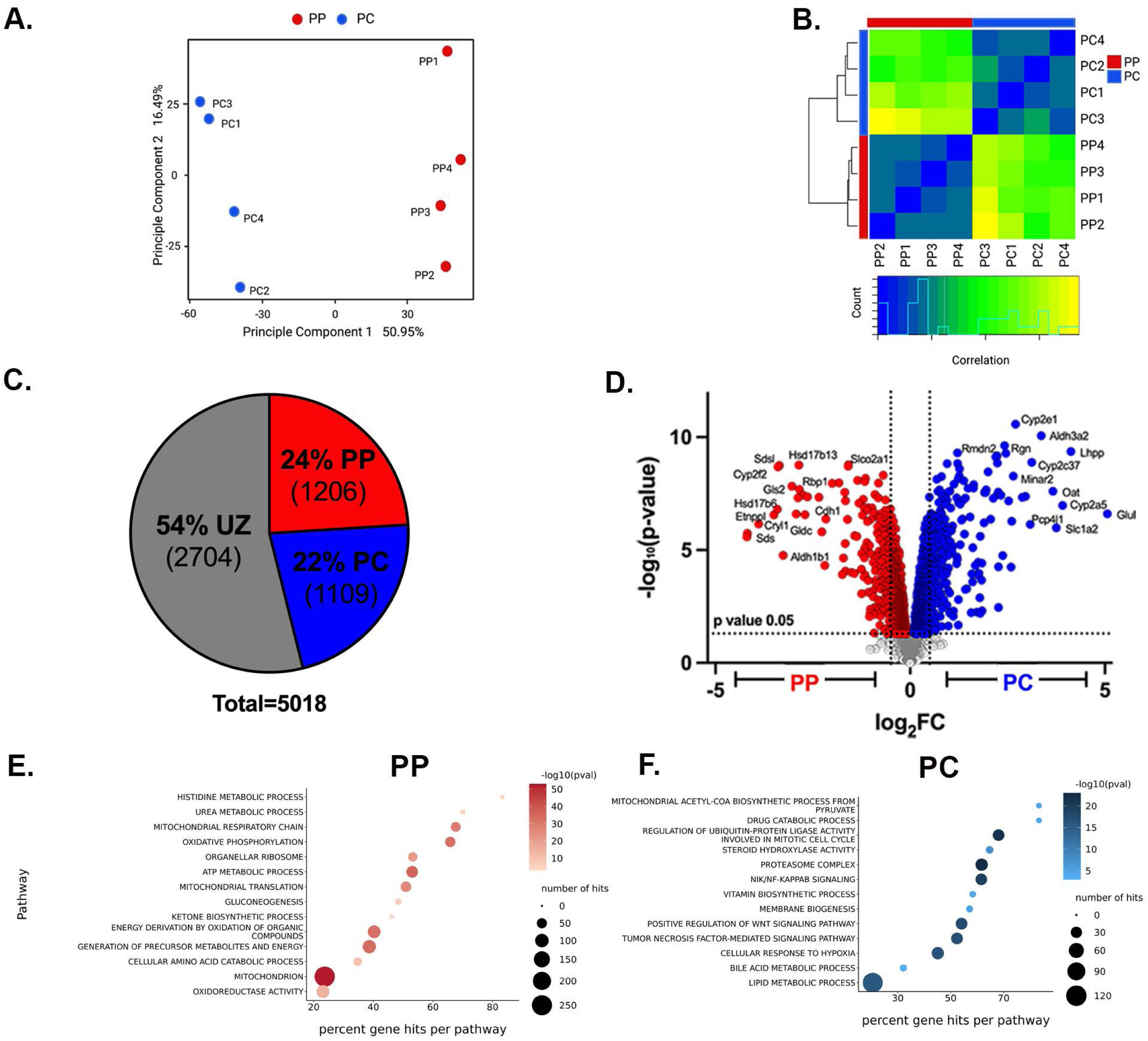
Comparative proteomics of spatially sorted hepatocytes. **(A)** Principal component analysis (PCA) and frequency histogram of spatially sorted hepatocytes analyzed with mass spectrometry. **(B)** Correlation Matrix Heatmap of proteomics data. **(C)** Pie chart depicting the percentage of PP, PC, and UZ proteins based on p-value (0.05). **(D)** Volcano plot showing the *PC* to PP log_2_ fold-change (x-axis) and the −log_10_ p-value (y-axis) for identified proteins. **(E and F)** GO enrichment analysis of the proteomics data in the spatially sorted cell.

**Figure S2.**
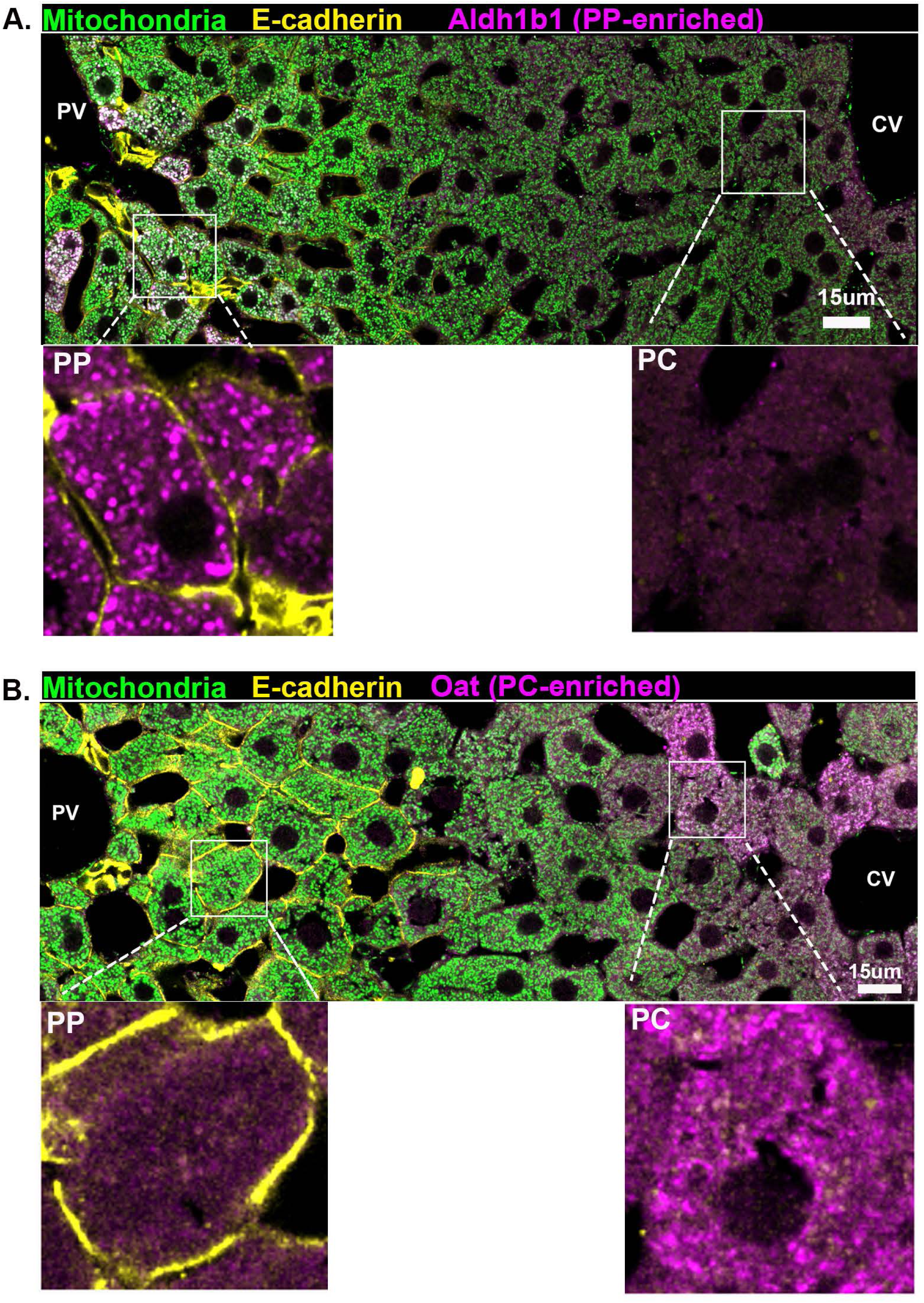
Immunofluorescence of representative PP and PC mitochondrial proteins. **(A)** Confocal image of the liver lobule from Mito-Dendra2 mice (green) stained with E-cadherin to label PP regions (yellow) and Aldh1b1, a PP mitochondrial protein (magenta). Magnified insets show the overlay of E-cadherin and Aldh1b1 only. Scale bar: 15 µm. **(B)** Confocal image of the liver lobule from Mito-Dendra2 mice (green) stained with E-cadherin to label PP regions (yellow) and Oat, a PC mitochondrial protein (magenta). Magnified insets show the overlay of E-cadherin and Oat only. Scale bar: 15 µm.

**Figure S3:**
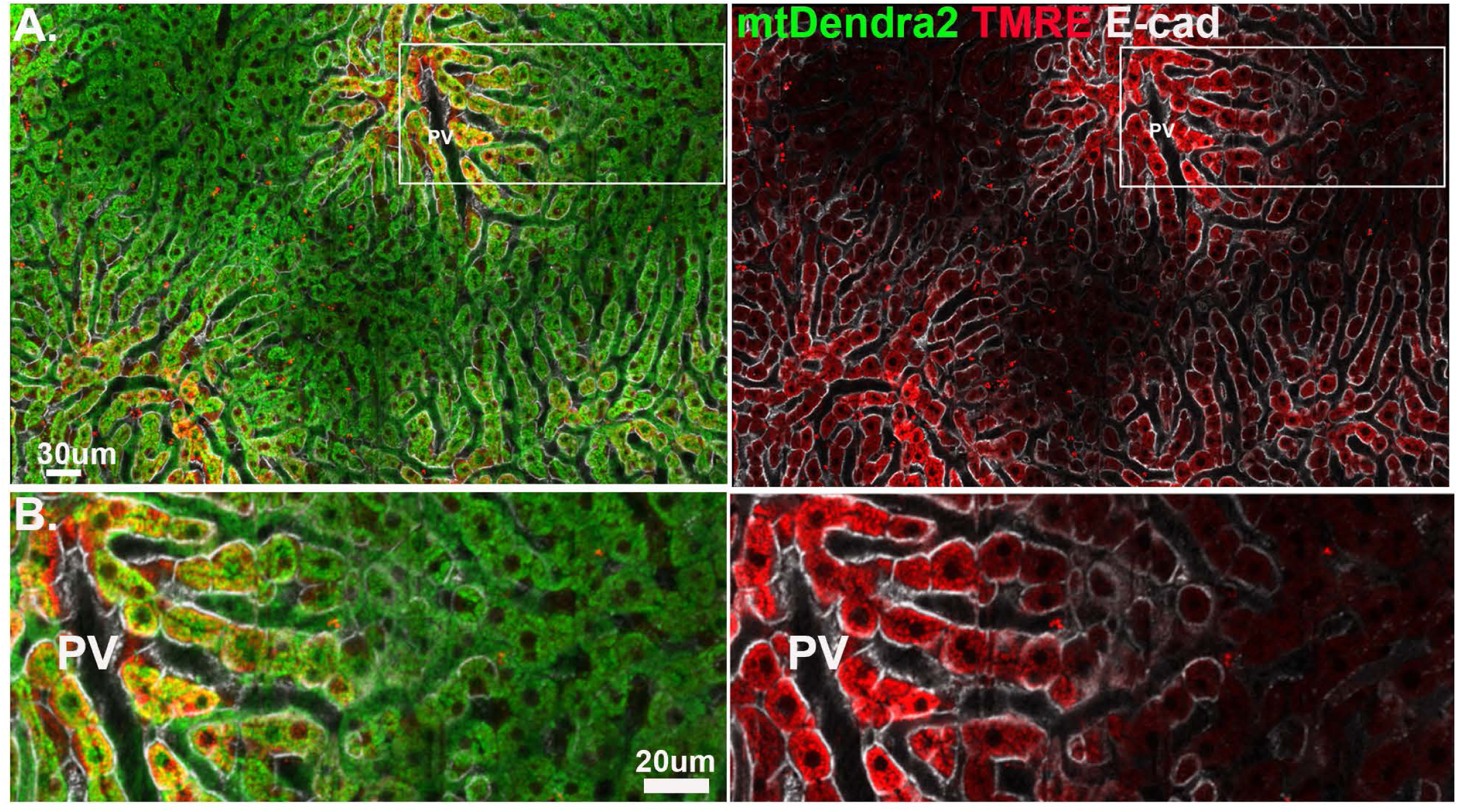
E-cadherin-positive hepatocytes display higher membrane potential. **(A-B)** Intravital microscopy of the hepatic lobule in Mito-Dendra2 mouse labeled with TMRE (red), and E-cadherin (white). **(A)** Low magnification of the hepatic lobule. Scale bar: 30 μm. **(B)** Close up on the PP-PC axis. Scale bar: 20 μm.

**Figure S4.**
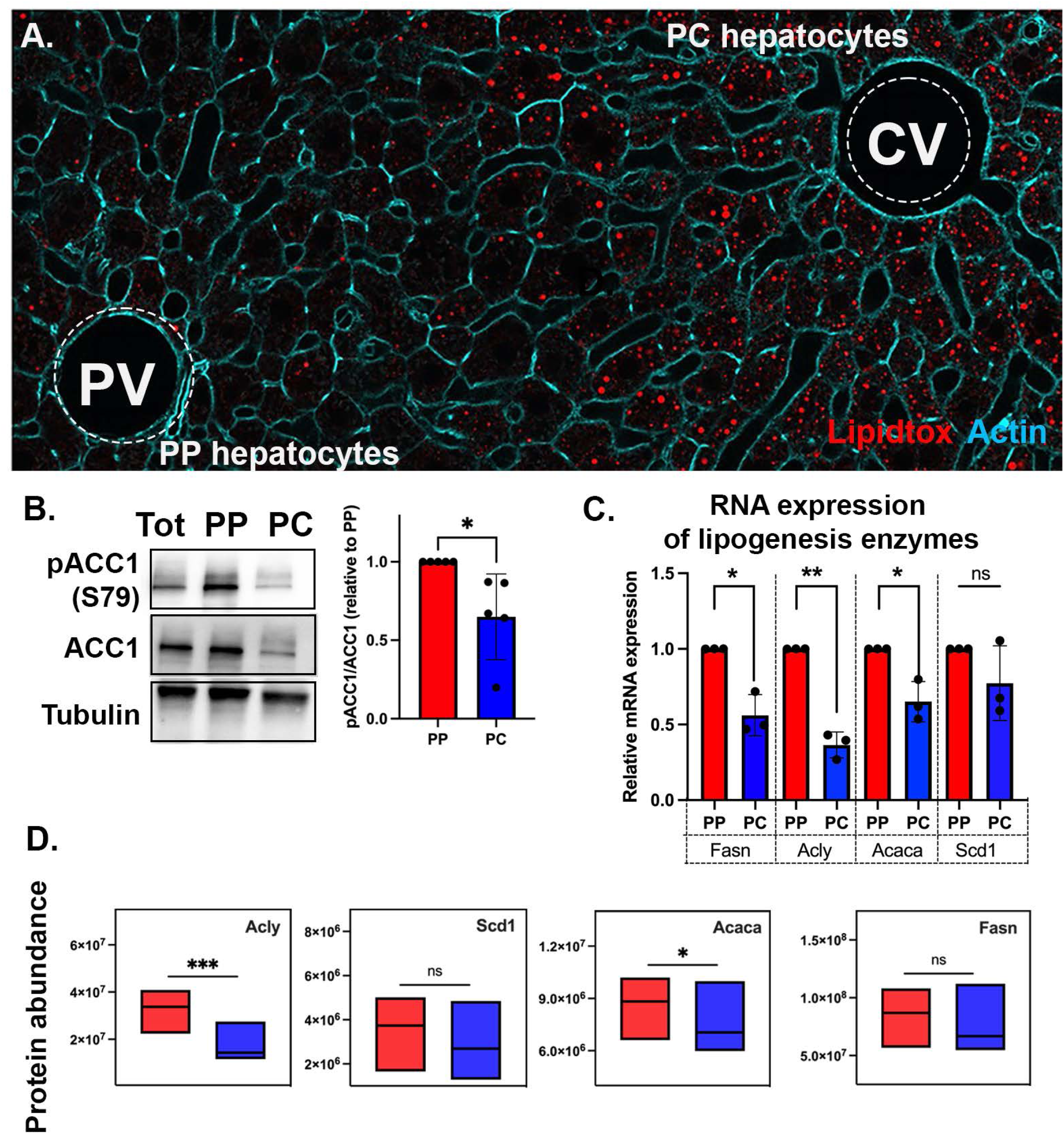
Lipid synthesis is pericentral in the murine liver. **(A)** Confocal image of a liver section labeled with Lipidtox (lipid droplets; red) and phalloidin (actin; cyan), showing the non-uniform distribution of lipid droplets across the PP-PC axis. **(B)** Representative immunoblot and quantification of acetyl-CoA carboxylase 1 (ACC1, Acaca) relative phosphorylation (S79) in spatially sorted hepatocytes. Bar graph shows five independent experiments. **(C)** RNA levels of key lipogenesis enzymes in spatially sorted hepatocytes. Bar graph shows three independent experiments. **(D)** Protein abundance of key lipogenesis enzymes identified by proteomics. Data presented as mean⍰±⍰SD. ns, **p <0.05, **p < 0.01, ***p⍰<⍰0.001.

**Figure S5.**
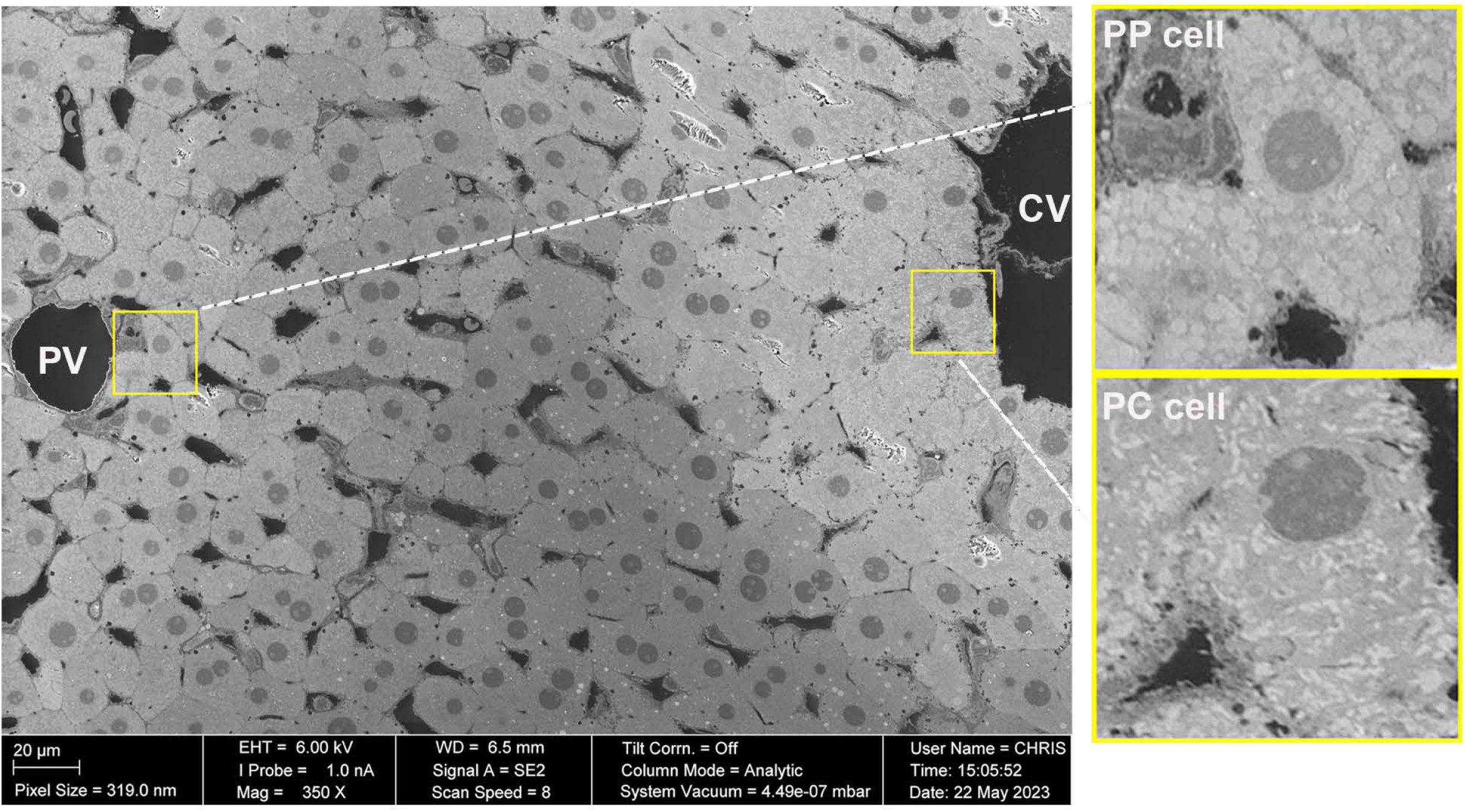
Scanning Electron Microscopy (SEM) of the lobule highlighting regions of interest selected for FIB-SEM. SEM of the PP-PC axis in a liver section was performed to mark PP and PC regions for FIB-SEM. Insets show PP and PC hepatocytes where striking variations in mitochondrial morphology are observed.

**Figure S6.**
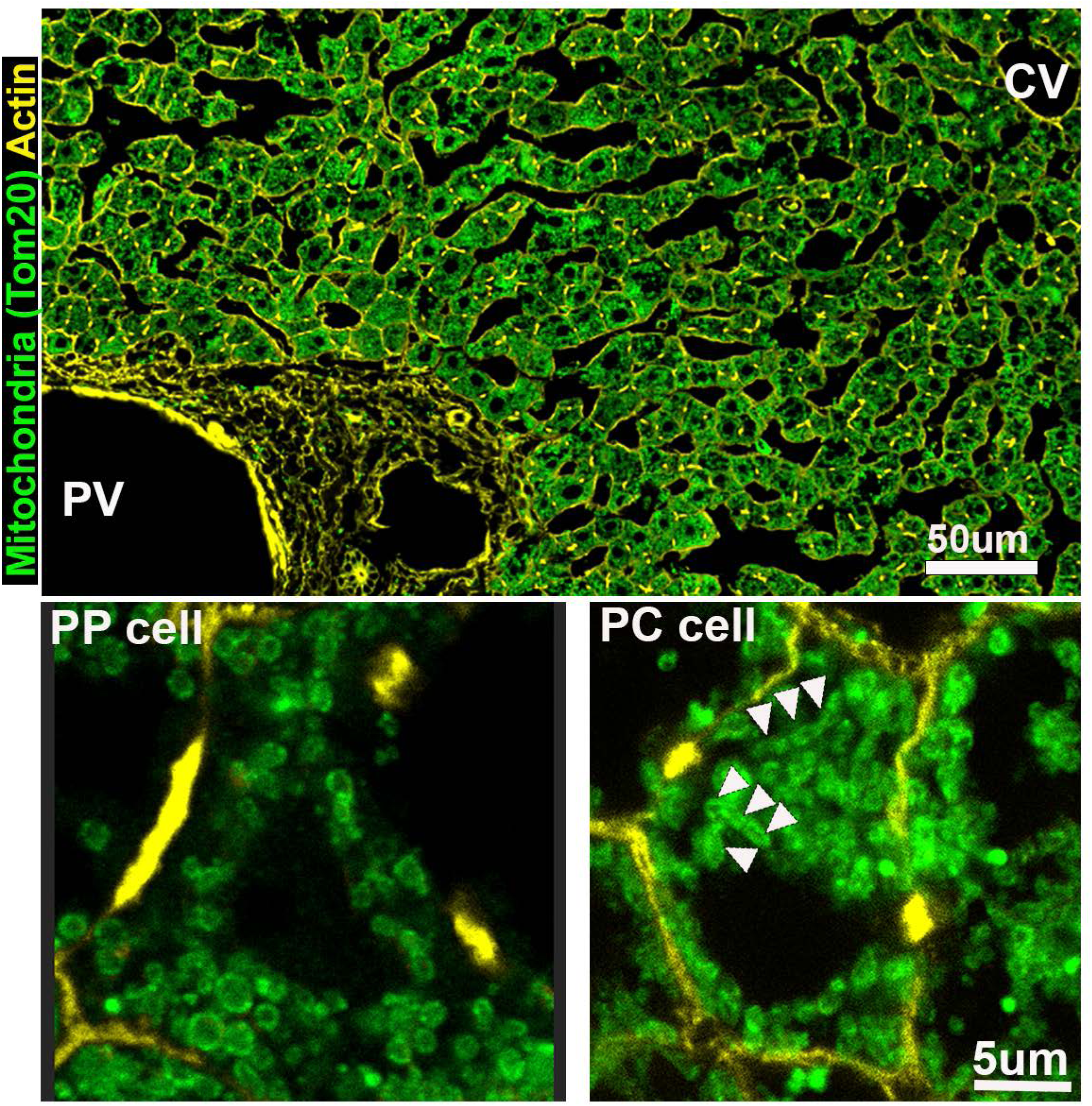
Mitochondrial morphologies are conserved in the human liver. Confocal image of fixed human liver sections labeled with phalloidin (actin; yellow) and TOM20 (outer mitochondrial membrane; green). Scale bar: 50μm. Insets of representative PP and PC hepatocytes show distinct morphological features (arrows). Scale bar: 5μm.

**Figure S7.**
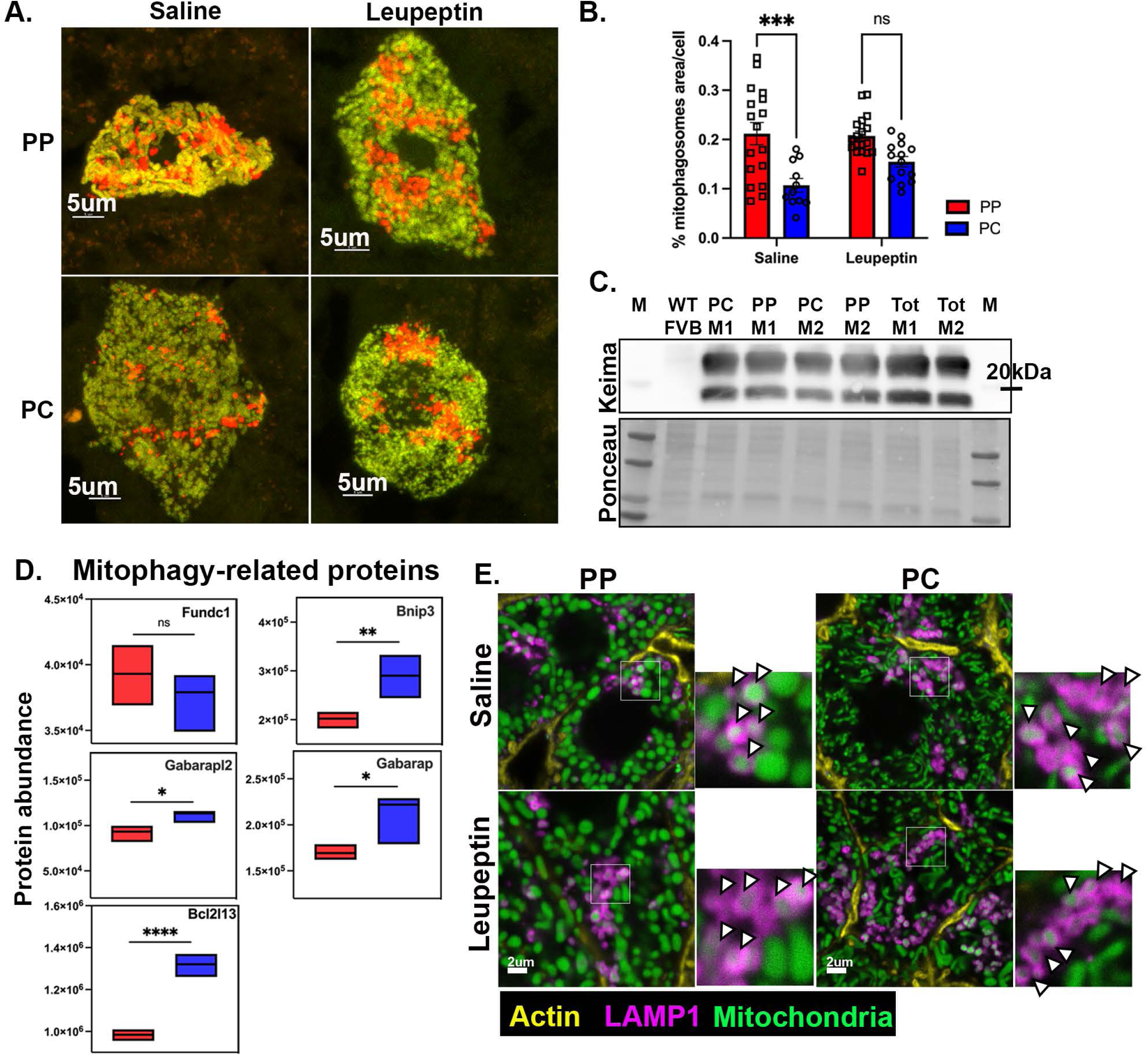
PC mitochondria display higher mitophagy flux. **(A)** Confocal microscopy of liver sections from mice transduced with Cox8-EGFP-mCherry adenovirus. Representative PP and PC cells are shown from saline or leupeptin-treated mice. Scale bar: 5μm. **(B)** The Cox8-EGFP-mCherry reporter was used to measure mitophagy by calculating the percent of the mitophagosome area normalized to the whole cell area in saline or leupeptin-treated mice. Bar graph shows 15-20 cells quantified from 3 mice. **(C)** Keima protein expression in hepatocytes isolated from one wild-type mouse and two mtKeima mice (M1 and M2). Sorted (PP and PC) and total are shown for mtKeima only. **(D)** The abundance of mitophagy-related proteins identified by proteomics is shown in box and whisker plots. Mitochondrial fission regulator 1 (Mtfr1); Bcl2 interacting protein 3 (Bnip3); GABA type A receptor-associated protein-like 2 (Gabarapl2); GABA type A receptor-associated protein like (Gabarapl). **(E)** Confocal images of PP and PC hepatocytes from Mito-Dendra2 mice (green), treated with saline or leupeptin and stained with the lysosomal marker LAMP1 (magenta) and actin (yellow). Arrowheads point to mitochondria (green) inside lysosomes (magenta). Data presented as mean⍰±⍰SD. ns, *p < 0.05, **p<0.01, ***p<0.001***p < 0.001.

**Figure S8.**
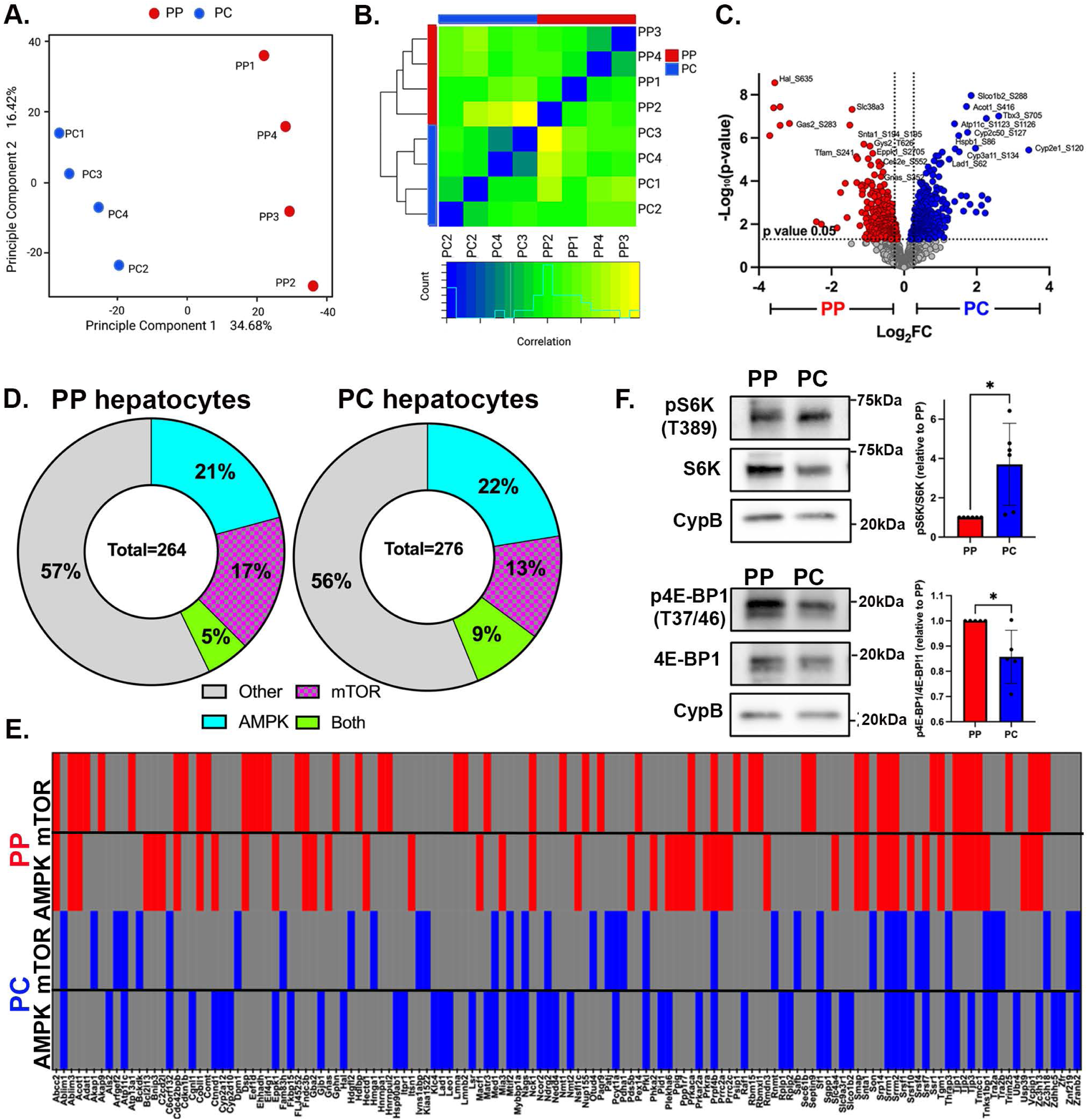
Comparative phosphoproteome in spatially sorted hepatocytes. **(A)** Principal component analysis (PCA) and **(B)** Correlation Matrix Heatmap of the phosphoproteome data set. **(C)** Volcano plot showing the log_2_ *PC/*PP fold-change (x-axis) and the −log_10_ p-value (y-axis). **(D)** Group-based Prediction System (GPS 5.0) was used to predict mTOR and AMPK phosphorylation sequences. **(E)** Complete list of mTOR and AMPK substrates identified in PP and PC. **(E)** Representative immunoblot and quantification of mTOR substrates pS6K (T389) and p4E-BP (T37/46) in spatially sorted hepatocytes. Bar graphs show five to six independent experiments. Data presented as mean⍰±⍰SD. ns, *p < 0.05.

**Figure S9.**
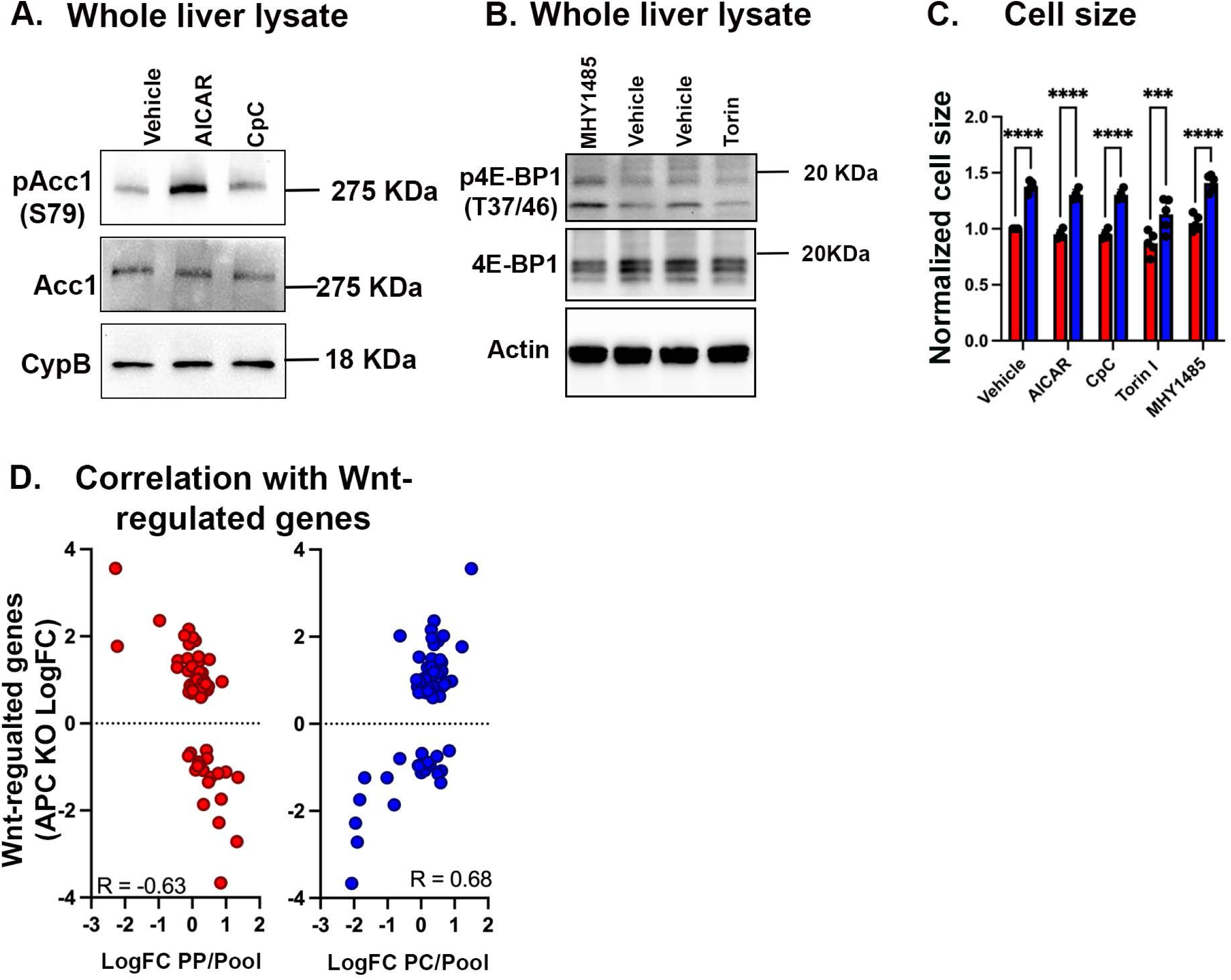
Nutrient sensing signaling regulates mitochondrial heterogeneity. **(A)** Determination of AICAR and Compound C (CpC) concentrations for *in vivo* modulation of AMPK signaling. Drug efficacy was evaluated by Western blotting for pACC1 (S79) levels. **(B)** Determination of MHY1485 and Torin concentrations for *in vivo* modulation of mTOR signaling. Drug efficacy was evaluated by Western blotting for p4EBP1(Thr37/Thr46) levels. **(C)** Mice were treated with two doses of vehicle, AMPK inhibitor (Compound C) or activator (AICAR), or mTOR inhibitor (Torin I) or activator (MHY1485). Graphs show the evaluation of drug effect on cell size measured by flow cytometry from three to five independent experiments. **(D)** Correlation plot of Wnt-activated genes and PP or PC mitochondrial proteome. Data presented as mean⍰±⍰SD. ns, *p < 0.05, **p < 0.01, ***p⍰<⍰0.001, ****p⍰<⍰0.0001.

**Movie S1.** Confocal z-stack of PP mitochondria

**Movie S2.** Confocal z-stack of PC mitochondria

**Movie S3.** FIB-SEM and volume rendering of PP mitochondria

**Movie S4.** FIB-SEM and volume rendering of PC mitochondria

**Supplementary files 1-3 and 6 will become available when the manuscript is accepted for publication**

**Supplementary File 4:** Modification Motifs (MoMo) analysis

**Supplementary File 5:** Extended table of mitochondrial phosphoproteins

## Material and methods

### Animal experiments

Experiments were approved by the Institutional Animal Care and Use Committee of the National Cancer Institute and comply with the Guide for the Care and Use of Laboratory Animals (National Institutes of Health publication 86-23, revised 1985). All experiments were conducted on ad libitum-fed, eight-to-ten-week-old male mice; C57BL/6J (strain# 000664), Mito-Dendra2 (strain# 018397) ^24^ obtained from Jackson Laboratories and the mtKeima line were a gift from Dr. Toren Finkel ^31^.

### Intra-cardiac fixation, tissue processing, and immunofluorescence

Mice were anesthetized with 250 mg/kg Xylazine, and 50 mg/kg Ketamine (diluted in saline) injected intraperitoneally (i.p.). The liver was fixed by transcardial perfusion of ice-cold PBS for 2 min followed by ice-cold 4% paraformaldehyde (PFA) in PBS at a rate of 5 ml/min. Livers were harvested and stored in 4% PFA in PBS overnight and processed in a sucrose gradient before embedding in OCT (Tissue-Tek). Blocks were kept at −80 °C until 10 μm thick slices were made with a cryostat and slides prepared. Slides were stored at −80 °C until thawed, rehydrated, and blocked with 0.1% Triton X-100 and 10% FBS in PBS for 1 h at room temperature. Next, slides were incubated with primary antibody at 4 °C overnight. The following day, slides were washed three times, 15 min each, then incubated with a secondary antibody for 1 h at room temperature. After three 15-min washes, slides were mounted with Fluoromount-G and a coverslip (#1).

### Confocal microscopy and image processing

Tile scans and z stacks were acquired using a Leica SP8 inverted confocal laser scanning microscope using a 63x oil objective and a 1.4 numerical aperture. Images were deconvolved using the LIGHTNING module in LAS X.

### Human tissue collection for immunofluorescence and imaging

Human tissue was obtained under an NIH IRB-approved protocol (13-C-0076) for risk-reducing surgery performed on patients with germline genetic mutation(s). All tissues procured, which included liver samples used in this study, were grossly normal as determined by the surgeon and histopathologically normal as determined by a board-certified pathologist. Of note, all tissue was obtained within 20 min of incision. Tissues were fixed overnight in 4% PFA in PBS overnight and processed in a sucrose gradient before embedding in OCT (Tissue-Tek).

### Measurement of mitochondrial membrane potential via intravital microscopy

Intravital microscopy was performed as previously described ^48^. Mitotracker Green FM (250 μM) and tetramethylrhodamine ethyl ester (TMRE, 200 μM) were consecutively injected retro-orbitally to label mitochondria with/without mitochondrial membrane potential in livers of C57BL/6J. After 30 min of incubation per dye, the liver was exposed to the microscope stage with the mouse under anesthesia, and images were acquired using excitation at 480 nm and 532 nm. The fluorescence intensity of TMRE and Mitotracker Green FM was analyzed across the PP-PC axis through line scanning with ImageJ. Additionally, Alexa Fluor® 647 anti-mouse/human CD324 (E-cadherin, 0.5 µg/g) was retro-orbitally injected to label PP regions in the liver of Mito-Dendra2 mice. Mito-Dendra2 mice were used instead of using Mitotracker Green FM dye to observe all mitochondria despite mitochondrial membrane potential and only TMRE (200 μM) was retro-orbitally injected and incubated for 30 min before imaging. Images were acquired using excitation at 480 nm, 532 nm, and 647 nm.

### Measurements of mitophagy

Mitophagy was measured in mtKeima mice ^31^ or C57BL/6J mice injected with Ad-Cox8-EGFP-mCherry (1×10^9^ PFU diluted in 200 μl of PBS) ^32^ using intravital microscopy as previously described ^48^. Lysosome inhibitor leupeptin (80 mg/kg), or saline was injected i.p. 16 h prior to the experiment. A second dose of leupeptin (40 mg/kg) or saline was administered 12 h later. Three to five mice were analyzed per experiment. This leupeptin treatment protocol was used prior to isolating and sorting cells into PP and PC populations for immunoblot analysis of mitophagy proteins.

### Western blot

Proteins from primary hepatocytes or whole liver were extracted by homogenization in RIPA lysis buffer (150 mM NaCl, 0.1% Triton X-100, 0.5% sodium deoxycholate, 0.1% sodium dodecyl sulfate, and 50 mM Tris HCl pH 8.0) containing EDTA, PMSF, and Halt Inhibitor Cocktail (Thermo Fisher), followed by centrifugation at 13,000 rpm at 4 °C for 30 min. Protein concentration was determined using a Pierce™ BCA Protein Assay Kit (Thermo Fisher). Lysates were heated at 95⍰°C for 5⍰min, and 5-10 μg aliquots fractionated by SDS-PAGE then transferred to nitrocellulose (0.45 μm) membrane (Bio-Rad). Membranes were blocked for 1 h at room temperature in 5% BSA in 1x Tris-buffered saline + 0.1% Tween 20 (TBST), then incubated with primary antibody diluted in 5% BSA in TBST at 4 °C overnight. Membranes were washed three times with TBST then incubated in secondary antibody 1 h at room temperature. Membranes were washed three times with TBST, then Clarity ECL western blot substrate solution (Bio-Rad) was applied for detection and imaged using the ChemiDoc Imaging System (Bio-Rad). Bands were quantified with ImageJ.

### Liver Perfusion and hepatocyte isolation

Livers of anesthetized mice were perfused by inserting a 22-gauge syringe into the portal vein and delivering 25 ml of pre-warmed (37 °C) perfusion buffer [Krebs-Henseleit buffer (Sigma) with 0.5 μM EDTA], followed by 25 ml collagenase A buffer [Krebs-Henseleit buffer, with 0.1 mM Ca_2_Cl and 0.4 mg/ml collagenase A (Sigma)]. After perfusion, livers were transferred to a Petri dish, flooded with cold PBS, and hepatocytes were gently released using forceps. Dissociated cells were collected and filtered through a 100 μm cell strainer. Cells were centrifuged at 100 G for 5 min at 4 °C to obtain hepatocyte-enriched cells. Pellets were resuspended in cold PBS, filtered through a 40 μm cell strainer, and centrifuged at 100 G for 5 min at 4 °C. Percoll diluted in 10x Hank’s buffer (Sigma) was added to the cell suspension and centrifuged at 100 G for 5 min at 4 °C. The supernatant containing dead cells was removed by aspiration. The pellet was resuspended in cold PBS, and cell number and viability were determined.

### Fluorescence-activated cell sorting (FACS) of PP and PC populations

Cell sorting was performed on a FACSAria Fusion cell sorter (BD Biosciences). Forward and side light scatter was used to distinguish cells from debris and to identify single cells. Zombie-Green fixable live dead dye (1:500, BioLegend) was used to discriminate between live and dead cells. PP and PC populations were identified using anti-E-cadherin-PE and anti-CD73-APC antibody staining (1:150; BioLegend). PP- and PC-specific gates were set after spectral compensation using appropriate fluorescent minus (FMO) controls. The protocol was developed based on ^49^.

To sort PP and PC hepatocyte populations from mtKeima mice, BD OptiBuild™ BUV395 Rat Anti-Mouse CD324/E-Cadherin (1:125, BD Bioscience) was used to stain PP hepatocyte populations. For E-cadherin negative populations (PC), FSC-A and SSC-A plot was used to further exclude smaller sized hepatocytes as PC hepatocytes (CD73 positive) was generally found to be bigger in cell size than PP hepatocytes.

### Protein Digestion and TMT labeling

Cells were lysed in 50 mM HEPES, pH 8.0, 8 M urea, and 10% methanol, followed by sonication. Lysates were clarified by centrifugation, and protein concentration quantified using a BCA protein estimation kit (Thermo Fisher). A 250 ug aliquot was alkylated and digested by incubating overnight at 37 °C in trypsin at a ratio of 1:50 (Promega). Digestion was acidified by adding formic acid (FA) to a final concentration of 1% and desalted using Pierce peptide desalting columns according to the manufacturer’s protocol. Peptides were eluted from the column using 50% ACN/0.1% FA, dried in a speed vac and kept frozen at −20 °C for further analysis.

For TMT labeling, 125 ug of each sample was reconstituted in 50 μl of 50 mM HEPES, pH 8.0, 500 ug of TMTpro-16plex label (Thermo Fisher) in 100% ACN. After incubating the mixture for 1 h at room temperature with occasional mixing, the reaction was terminated by adding 8 μl of 5% hydroxylamine. As there were more than 16 samples, we generated a pooled sample consisting of equal amounts of lysate from each condition and TMT labeled. The pool was added to each TMT experiment. The TMT-labeled peptides were pooled and dried in a speed vac. The samples were desalted, and excess TMT labels were removed using peptide desalting columns (Thermo Fisher). A 100 ug aliquot of labeled peptide mixture was fractionated using high pH reversed-phase, and the remaining peptide mixture was used for phospho-enrichment.

### Phosphopeptide enrichment

Phosphopeptides were enriched from the TMT-labeled peptides using the SMOAC (Sequential enrichment of Metal Oxide Affinity Chromatography) method. First, the phosphopeptides were enriched using the HiSelect TiO_2_ phosphopeptide enrichment kit (Thermo Fisher) according to the manufacturer’s protocol. The flow-through and wash from the TiO_2_ enrichment kit were then combined and enriched using the HiSelect Fe-NTA phosphopeptide enrichment kit (Thermo Fisher). The enriched phosphopeptides obtained by both methods were dried by speed vac and stored at −20 °C until analysis by mass spectrometry.

### High pH reversed-phase fractionation

The first-dimensional separation of peptides was performed using a Waters Acquity UPLC system coupled with a fluorescence detector (Waters) using a 150 mm x 3.0 mm Xbridge Peptide BEM^TM^ 2.5 μm C18 column (Waters) operating at 0.35 ml/min. The dried peptides were reconstituted in 100 μl of mobile phase A solvent (3 mM ammonium bicarbonate, pH 8.0). Mobile phase B was 100% acetonitrile (Thermo Fisher). The column was washed with mobile phase A for 10 min followed by gradient elution 0-50% B (10-60 min) and 50-75% B (60-70 min). Fractions were collected every minute. These 60 fractions were pooled into 24 fractions. Fractions were vacuum centrifuged, and lyophilized fractions were stored at −80°C until analysis by mass spectrometry.

### Mass Spectrometry acquisition

The lyophilized peptide fractions were reconstituted in 0.1% TFA and subjected to nanoflow liquid chromatography (Thermo Ultimate^TM^ 3000RSLC nano LC system, Thermo Fisher) coupled to an Orbitrap Eclipse mass spectrometer (Thermo Fisher). Peptides were separated using a low pH gradient and 5-50% ACN over 120 min in a mobile phase containing 0.1% formic acid at a 300 nl/min flow rate. MS scans were performed in the Orbitrap analyzer at a resolution of 120,000 with an ion accumulation target set at 4e^5^ and max IT set at 50 ms over a mass range of 400-1600 m/z. Ions with a determined charge state between 2 and 5 were selected for MS2 scans in the ion trap with CID fragmentation (Turbo; NCE 35%; maximum injection time 35⍰ms; AGC 1⍰×⍰10^4^). The spectra were searched using the Real Time Search Node in the tune file using mouse Uniprot database using Comet search algorithm with TMT16 plex (304.2071 Da) set as a static modification of lysine and the N-termini of the peptide. Carbamidomethylation of cysteine residues (+57.0214⍰Da) was set as a static modification, while oxidation of methionine residues (+15.9949⍰Da) was set up as a dynamic modification. For the selected peptide, an SPS–MS3 scan was performed using up to 10 *b*- and *y*-type fragment ions as precursors in an Orbitrap at 50,000 resolution with a normalized AGC set at 500 followed by maximum injection time set as “Auto” with a normalized collision energy setting of 65.

The enriched phospho-samples were reconstituted in 0.1% TFA and analyzed using an Orbitrap Eclipse mass spectrometer. Peptides were separated using a low pH gradient, and FAIMS was enabled during data acquisition with the compensation voltages set at −45V, −60V, and −75V set up at the first run, −50V, −65V, and −80V for the second run and −55V, −70V and −85V for the last run. MS scans were performed in the Orbitrap analyzer at a resolution of 120,000 with an ion accumulation target set at 4e^5^ and max IT set at 50ms over a mass range of 350-1600 m/z. Ions with a known charge state between 2 and 6 were selected for MS2 scans. A cycle time of 1 sec was used for each CV, and a quadrupole isolation window of 0.4 m/z was used for MS/MS analysis. An Orbitrap at 15,000 resolution with a normalized AGC set at 250 followed by maximum injection time set as “Auto” with a normalized collision energy setting of 38 was used for MS/MS analysis. The node “Turbo TMT” was switched on for the high-resolution acquisition of TMT reporter ions.

## Data Analysis

Acquired MS/MS spectra were searched against the mouse Uniprot protein database and a contaminant protein database using SEQUEST in the Proteome Discoverer 2.4 software (Thermo Fisher). The precursor ion tolerance was set at 10 ppm, and the fragment ions tolerance was set at 0.02 Da, with methionine oxidation included as a dynamic modification. For the phospho-enriched samples, serine, threonine, and tyrosine phosphorylation were set as variable modifications. Carbamidomethylation of cysteine residues and TMTpro16 plex (304.2071 Da) was set as static modification of lysine and the N-terminus of the peptide. Trypsin was specified as the proteolytic enzyme, with up to 2 missed cleavage sites allowed. Searches used a reverse sequence decoy strategy to control for the false peptide discovery, and identifications were validated using the Percolator algorithm in Proteome Discoverer 2.4.

Reporter ion intensities were adjusted to correct for lot-specific impurities according to the manufacturer specification, and the abundances of the proteins were quantified using the summation of the reporter ions for all identified peptides. A two-step normalization procedure was applied to a 24-plex TMT experiment [2 TMT experiments with 12 samples each]. First, the reporter abundances were normalized across all the channels to account for equal peptide loading. Secondly, the intensity from the pooled sample was used to normalize the batch effect from the multiple TMT experiments. Samples were further normalized using the Voom algorithms and quantile normalization from the Limma R package (v3.40.6). We evaluated the significance of a given protein, phosphopeptide, or normalized phosphopeptide change using Limma, where zonation was determined by the p-value (0.05) and/or FC (³1.2). Pathway enrichment analysis was performed using Fisher’s Exact test with GO database. MitoCarta 3.0 was used to classify of the identified proteins and phosphoproteins into mitochondrial and non-mitochondrial proteins. GraphPad Prism 9 and BioRender were used for visualization.

### Prediction of mTOR and AMPK activity using Group-based Prediction System (GPS)

All peptides with PP or PC zonation were extrapolated as 15 AA sequence with the phosphosite in the center. Peptides without a specific site were excluded. Duplicate sequences were removed, and sequences were analyzed using GPS 5.0 (AMPK and mTOR selected) at medium threshold. The sequences analyzed and the output from GPS are listed in Supplementary File 6.

### MoMo motif analysis

Motif analyses of PP or PC phosphopeptides were performed using the web tool available at: https://meme-suite.org/meme/tools/momo, v5.5.4. All identified 15 AA sequences without duplicates for each PP and PC quantified phosphosite were analyzed with a p-value threshold at 10^-6^. The results from this analysis are listed in Supplementary File 4.

### WNT-activated genes correlation

We examined identified mitochondrial proteins associated with WNT signaling transduction^50^. PP and PC were separately analyzed by computing the log_2_ ratio of the mean protein abundance over the geomean from the raw data. These values were then used to examine Spearman correlation value with log2 ratio of the mean gene expression levels from WNT-hyperactivating liver-specific APC knockout and control mouse liver samples.

### Seahorse assay

A Seahorse XFe96 Analyzer and XF Mito Stress Test Kit (Agilent Technologies) were used to measure the oxygen consumption rate (OCR) and extracellular acidification rate (ECAR) of PP and PC-sorted cells. A day before the assay, the cartridge sensor was hydrated overnight with Seahorse Bioscience Calibration Solution at 37⍰°C without CO_2_. After sorting, cells were seeded at 5.0⍰×⍰10^4^cells/well, in a 96-well seahorse plate and allowed to adhere for 2 h. After 2 h, media was replaced with serum-free Seahorse XF Base media containing glucose, pyruvate, and glutamine (pH⍰7.4), and cells were incubated at 37 °C in a non-CO_2_ incubator for 1 h. OCR and ECAR were measured after the injection of oligomycin (1.5 μM), FCCP (1 μM), and rotenone/antimycin (0.5 μM). The XF Mito Stress Test Kit was used together with an XF Mito Fuel kit to test fuel dependency. Three inhibitors, UK5099 (2 μM), BPTES (3 μM), and etomoxir (4 μM), were added to measure the dependency of cells on the three primary mitochondrial fuels to produce ATP; glucose, glutamine, and long-chain fatty acids, respectively. Average protein amount of PP and PC seeded wells were examined with BCA for normalization. To overcome reading variations within each population, and measure the effect of the inhibitors, the measurements of the wells treated with inhibitors were further normalized to the first basal reading of the group treated with the vehicle (control). Measurements for ATP production rate and maximum respiration capacity were presented as a percent normalized to PP. Bar graph shows data from four independent experiments presented as mean±SD. Each independent experiment was conducted with five to twelve replicates per population and treatment.

### ATP measurement assay

The bioluminescence ATP assay kit was used according to manufacturer instructions (Invitrogen) to analyze spatially sorted hepatocytes. ATP levels were calculated and normalized according to protein amount.

### Citrate Synthase activity assay

Citrate Synthase (CS) activity was analyzed following the manufacturer protocol (Abcam). After collecting the cells via FACS, cells were lysed. A 50 μl aliquot of supernatant or standard GSH solution with 50 μl of reaction buffer was added per well in a 96-well plate to measure the absorbance (OD 412 nm) in kinetic mode at 25 °C for 40 min. CS activity was calculated based on two different time points considering the dilution factor and cell number, then normalized to the PP fraction.

### Triglyceride (TG) measurement assay

After cell isolation and FACS, the sorted PP and PC cells were pelleted, washed twice with cold PBS, and lysed on ice for 30 min. Lysates were centrifuged at 13,000 rpm for 10 min at 4 °C and the supernatant collected. A TG-colorimetric GPO-PAP assay kit (Elab Science) was used to determine TG content, following manufacturer instructions. TG levels were calculated according to protein amount and normalized to PP fraction.

### DNA and RNA isolation

RNA was isolated from spatially sorted cells using the RNeasy mini kit and QIAshredder homogenizer columns (Qiagen). Total DNA was isolated using the DNeasy Blood and Tissue kit (Qiagen).

### Quantitative PCR

A High-Capacity RNA to cDNA kit (Thermo Fisher) was used to synthesize random-primed cDNA from 600 ng DNAse treated RNA. Real-time PCR was conducted in 384 well-plates using a ViiA7 Real-time PCR system (Applied Biosystems). Singleplex reactions (10 μl) containing a FAM-MGB expression assay for the gene of interest (Acaca Mm01304289_m1, Acly Mm01302282_m1, Fasn Mm00662319_m1, Scd1 Mm00772290_m1) or endogenous control (Ppia Mm02342430_g1) (Thermo Fisher) were performed using cDNA synthesized from 3 ng RNA and 1x Fast Advanced Master Mix (Thermo-without Amp Erase UNG). The comparative Ct method (delta, delta Ct) was used to determine relative expression normalized to Ppia (Applied Biosystems® ViiA™ 7Real-Time PCR System Getting Started Guides).

### mtDNA copy number determination

Mitochondrial DNA content was measured by comparing the DNA copy number of two mitochondrial encoded genes (ND1 and 16S rRNA) to HK, a nuclear-encoded gene^51^. The mitochondrial to nuclear DNA ratio (mtDNA/nDNA) was determined by Real-time PCR using an ABI Viia7 Real-Time PCR System (Applied Biosystems). Because mitochondrial genes lack introns, it was possible to use TaqMan gene expression assays for mtDNA copy number. Singleplex reactions (10 μl) containing FAM-MGB assays for ND1 Mm04225274_s1, Rnr2 Mm04260181_s1, or HK2 Mm00193901_cn (Applied Biosystems) were performed in quadruplicate using 384 well plates with 10 ng DNA and 1x Universal Master Mix (Applied Biosystems-without Amp Erase UNG). Determination of mtDNA content was performed using the comparative Ct method (delta, delta Ct) with normalization to HK2 as an internal reference (Applied Biosystems® ViiA™ 7Real-Time PCR System Getting Started Guides).

### Drug treatment and flow cytometry assay

Mice were treated with two doses of either vehicle, Compound C (CpC, 20 mg/kg), 5-Aminoimidazole-4-carboxamide ribonucleotide (AICAR, 0.5 mg/kg), Torin I (20 mg/kg) or MHY1485 (20 mg/kg) before primary hepatocytes were isolated via perfusion. Hepatocytes were then labeled with Cy-PE/PE-anti-E-cadherin and APC-anti-CD73 and either JC1 (5 μM) or BODIPY-493/503 (2 μM) for 20 min at 37 °C. The fluorescent signal was measured by BD LSRFortessa (BD Biosciences) cell analyzer. Consistent with FACS, forward and side light scatter was used to distinguish cells from debris and to identify single cells.

### Sample prep and imaging for FIB-SEM

The structured sample preparation protocol based on the sample preparation widget from EMPIAR (https://www.ebi.ac.uk/empiar/widget) can be accessed at: 10.5281/zenodo.8422314. Briefly, lower left lobes dissected from normally fed mouse livers were fixed in Karnovsky’s fixative for 2 h at room temperature, rinsed (all rinses were typically five times 3 min each at room temperature) in 0.1 M sodium cacodylate buffer, then fixed in 2% w/v osmium tetroxide + 1.5% w/v potassium ferricyanide in sodium cacodylate for 1 h at room temperature. The biosamples were then rinsed in dd-water, then *en bloc* stained with 1% uranyl acetate in dd-water overnight at 4 °C. These were then rinsed in dd-water and incubated in freshly made lead aspartate for 30 min at 60 °C. After another dd-water rinse, the biosamples were dehydrated for 10 min each in 35%, 50%, 70%, 95%, three times in 100% ethanol, and finally in three times in 100% propylene oxide. The dehydrated tissues were infiltrated with Polybed 812 resin (hard formulation) in resin: PO ratios of 1:3 (1 h), 1:1 (overnight), 3:1 (5 h) and finally 100% resin overnight. These were transferred to beam capsules and the resin was cured at 60 °C for 48 h to yield samples ready for vEM. Liver samples were sectioned and stained with Toluidine Blue to identify periportal and pericentral regions in the tissue. The remaining sample was trimmed and mounted on an SEM stub, and after a thin coat (approximately 8 nm) of carbon was sputtered, the specimens were into a FIB-SEM (Zeiss crossbeam 550). The same areas were located using SEM imaging, and Fields of View (FOVs) for vEM imaging were defined. Once the target cell was identified, the Fibics Atlas 3D FIB sample preparation workflow was initiated. A 1um thick protective platinum pad was deposited over the sample using a FIB current of 1.5 nA, after which tracking and autofocus lines were milled into the platinum surface and subsequently covered by a FIB mediated deposition of 1 um carbon at 1.5 nA. A coarse trench was milled using a 30nA FIB beam and fine polished using a 3nA beam. Milling and imaging parameters during the “continuous mill-and-image” acquisition run were set at 30kV accelerating voltage, 1.5nA FIB current, and 1.5kV accelerating voltage, 1.1nA SEM beam current. A pixel resolution of 5 nm with a 10 nm slice thickness was set, with total dwell time of 3.0 us/pixel, and the EsB detector grid voltage was set at 825 V. Image frame time was approximately 42 s, FIB advance rate was approximately 13.2 nm/min. The raw image stacks were then registered, inverted and binned using in-house scripts ^52^.

### Mitochondria segmentation and image analysis

Volume EM 3D reconstructions were imported into napari (https://napari.org), and mitochondria were segmented using the empanada plugin (https://empanada.readthedocs.io/en/latest/empanada-napari.html) by deploying the deep learning model MitoNet-mini with standard presets and ortho-inference^53^. The model produced high-quality mitochondrial segmentations, with some split and merge errors as expected for crowded mitochondria. These errors were manually corrected using empanada to produce instance segmentations of mitochondria from PP and PC volumes. Mitochondria that were partially cut off by the image limits were computationally removed to prevent artifactual measurements. The surface area, volume, and sphericity of remaining mitochondria (PP: n=175, PC: n=250) were measured for each instance using in-house scripts based on the scikit-image library (https://scikit-image.org/).

