## Supplementary material for "A spatial map of hepatic mitochondria uncovers functional heterogeneity shaped by nutrient-sensing signaling": Sup File 4

| mod | motif | regexp | score | fg_match | fg_size | bg_match | bg_size | fg/bg | unadjusted_p-value | tests | adjusted_p-value |
| --- | --- | --- | --- | --- | --- | --- | --- | --- | --- | --- | --- |
| S | xxxxxxx_S_PxPxxxx | .......SP.P.... | 45.6 | 23 | 220 | 1 | 220 | 23 | 8.50E-07 | 301 | 2.60E-04 |
| S | xxxxxxx_S_Pxxxxxx | .......SP...... | 22.33 | 88 | 220 | 19 | 220 | 4.6 | 3.40E-15 | 257 | 8.80E-13 |
| S | xxxxxxx_S_DxExxxx | .......SD.E.... | 14.39 | 12 | 220 | 1 | 220 | 12 | 1.50E-03 | 178 | 2.30E-01 |
| S | xxxxRxx_S_xxxxxxx | ....R..S....... | 6.62 | 61 | 220 | 34 | 220 | 1.8 | 1.20E-03 | 180 | 2.00E-01 |
| T | xxxxxxx_T_Pxxxxxx | .......TP...... | 9.12 | 20 | 28 | 5 | 28 | 4 | 5.90E-05 | 23 | 1.40E-03 |

PP

PC

| mod | motif | regexp | score | fg_match | fg_size | bg_match | bg_size | fg/bg | unadjusted_p-value | tests | adjusted_p-value |
| --- | --- | --- | --- | --- | --- | --- | --- | --- | --- | --- | --- |
| S | xxxxxSx_S_Pxxxxxx | .....S.SP...... | 33.34 | 22 | 238 | 1 | 238 | 22 | 1.80E-06 | 265 | 4.80E-04 |
| S | xxxxxxx_S_Pxxxxxx | .......SP...... | 11.48 | 68 | 238 | 17 | 238 | 4 | 4.00E-10 | 222 | 8.90E-08 |
| S | Kxxxxxx_S_xxxxxxx | K......S....... | 8.01 | 25 | 238 | 7 | 238 | 3.6 | 7.40E-04 | 172 | 1.20E-01 |
| S | xxxxRxx_S_xxxxxxx | ....R..S....... | 7.44 | 69 | 238 | 35 | 238 | 2 | 1.10E-04 | 191 | 2.20E-02 |
| S | xxxxxxx_S_Dxxxxxx | .......SD...... | 8.29 | 37 | 238 | 19 | 238 | 1.9 | 7.50E-03 | 135 | 6.40E-01 |
| T | xxxxxxx_T_Pxxxxxx | .......TP...... | 7.82 | 17 | 27 | 4 | 27 | 4.2 | 3.10E-04 | 21 | 6.50E-03 |
