## Supplementary material for "A spatial map of hepatic mitochondria uncovers functional heterogeneity shaped by nutrient-sensing signaling": Sup File 5

| **Protein** | **P-Zonation** | **P-site** | **Upstream regulator/kinase** | **Function** |
| --- | --- | --- | --- | --- |
| Acaca | PP | S29 | Leptin, Raptor, BMP2, AMPK(1) | Lipogenesis |
| Akap1 | PP | S101/S103/S104/S109 | ADRB1, RIPK3  Insulin, LPA, LY294002, AMPK(2) | cAMP-PKA signaling |
| Apex1 | PP | S18 |  | mtDNA maintenance |
| Bcl2l13 | PP | S387 | Leptin | Mitophagy |
| Bnip3 | PP | S79  S85  S88  T66 | TSC2  ADRB1, TSC2  ADRB1, Rictor, STK3/4(3)  JNK1/2(4) | Mitophagy |
| Comt | PP | S261 | Refeeding | Catechol metabolism |
| Ehhadh | PP | T543 | Leptin | Fatty acid oxidation |
| Gpam | PP | S687  S694 | Refeeding  Leptin, Refeeding | Phospholipid metabolism |
| Mtfr1 | PP | T118 |  | Fission |
| Mtfr1l | PP | S234/S235 | Leptin, AMPK(5) | Fission |
| Nadk2 | PP | S373 |  | NAD biosynthesis and metabolism |
| Prkaca | PP | T198 | PDK1(6), PKACA, ADRB1, GPR107, Fasting | cAMP-PKA signaling |
| Rmdn3 | PP | S46  S50 | Leptin, PTH(1-34), PKA(7)  Insulin | Mitochondrial dynamics/contact sites |
| Slirp | PP | S105 | Leptin | mtRNA stability |
| Tfam | PP | S241 |  | mtDNA maintenance |
| Tomm70 | PP | S94 | ADRB1, Leptin, CK2α(8) | Protein import and sorting |
| Akap1 | PC | T487 |  | cAMP-PKA signaling |
| Bckdk | PC | S31 | APN(9) | Amino acid metabolism |
| Bnip3 | PC | S60 | JNK1/2(4) | Mitophagy |
| Coq9 | PC | S81 | Leptin, BMP2, SB202190 | Coenzyme Q metabolism |
| Gpam | PC | S694 | Leptin, Refeeding | Phospholipid metabolism |
| Hsd17b8 | PC | S58 |  | Lipid biosynthesis |
| Kars1 | PC | S594 |  | Protein translation |
| Miga2 | PC | S276 |  | Fusion/contact sites |
| Mrps36 | PC | S55/T59/S60 |  | Mitochondrial translation |
| Mtif2 | PC | S180 |  | Protein translation |
| Nags | PC | S44 |  | Amino acid metabolism |
| Nsun2 | PC | S23 | Rictor | RNA modification |
| Pdha1 | PC | S293, S300 | PDK1/3/4(10), AMPKA2, AS184285, BCKDH E1α, DCA, Fasting, Hypoxia | Pyruvate metabolism |
| Tomm22 | PC | S45 | Insulin | Protein import and sorting |

| ADRB1 | Adrenoceptor Beta 1 |
| --- | --- |
| AMPK | AMP-activated protein kinase |
| AMPKA2 | Protein Kinase AMP-Activated Catalytic Subunit Alpha 2 |
| APN | Aminopeptidase N |
| AS184285 | Foxo1 (Forkhead Box O1) Inhibitor |
| BCKDH E1a | Branched Chain Keto Acid Dehydrogenase E1 Subunit Alpha |
| BMP2 | Bone Morphogenetic Protein 2 |
| CK2a | Casein Kinase 2 Alpha 2 |
| GPR107 | G Protein-Coupled Receptor 107 |
| JNK1/2 | c-Jun N-terminal kinase 1/2 |
| LPA | Lipoprotein(A) |
| LY294002 | Phosphoinositide 3-kinase inhibitor |
| PDK1-4 | Pyruvate Dehydrogenase Kinase 1-4 |
| PKA | Protein kinase A |
| PKACA | Protein Kinase CAMP-Activated Catalytic Subunit Alpha |
| PTH(1-34) | Selective activator of the parathyroid hormone receptor (PTH1R) signaling pathway |
| Raptor | Regulatory-associated protein of mTOR |
| Rictor | Rapamycin-insensitive companion of mammalian target of rapamycin |
| RIPK3 | Receptor Interacting Serine/Threonine Kinase 3 |
| SB202190 | MAPK inhibitor |
| STK3/4 | Serine/Threonine Kinase 3/4 |
| TSC2 | Tuberin |

Hornbeck PV, Zhang B, Murray B, Kornhauser JM, Latham V, Skrzypek E PhosphoSitePlus, 2014: mutations, PTMs and recalibrations. Nucleic Acids Res. 2015 43:D512-20

1. Activity and structure of human acetyl-CoA carboxylase targeted by a specific inhibitor

SoRi Jang, Piotr Gornicki, Jasmina Marjanovic, Ethan Bass, Toni P. Iurcotta, Pedro Rodriguez, Jotham Austin II, Robert Haselkorn

First published: 17 May 2018 https://doi.org/10.1002/1873-3468.13097

2. Global Phosphoproteomic Analysis of Human Skeletal Muscle Reveals a Network of Exercise-Regulated Kinases and AMPK Substrates.

Hoffman NJ, Parker BL, Chaudhuri R, Fisher-Wellman KH, Kleinert M, Humphrey SJ, Yang P, Holliday M, Trefely S, Fazakerley DJ, Stöckli J, Burchfield JG, Jensen TE, Jothi R, Kiens B, Wojtaszewski JF, Richter EA, James DE. Cell Metab. 2015 Nov 3;22(5):922-35. doi: 10.1016/j.cmet.2015.09.001. Epub 2015 Oct 1. PMID: 26437602

3. STK3/STK4 signalling in adipocytes regulates mitophagy and energy expenditure.

Cho YK, Son Y, Saha A, Kim D, Choi C, Kim M, Park JH, Im H, Han J, Kim K, Jung YS, Yun J, Bae EJ, Seong JK, Lee MO, Lee S, Granneman JG, Lee YH. Nat Metab. 2021 Mar;3(3):428-441. doi: 10.1038/s42255-021-00362-2. Epub 2021 Mar 23. PMID: 33758424

4. BNIP3 phosphorylation by JNK1/2 promotes mitophagy via enhancing its stability under hypoxia.

He YL, Li J, Gong SH, Cheng X, Zhao M, Cao Y, Zhao T, Zhao YQ, Fan M, Wu HT, Zhu LL, Wu LY. Cell Death Dis. 2022 Nov 17;13(11):966. doi: 10.1038/s41419-022-05418-z. PMID: 36396625

5. AMPK-dependent phosphorylation of MTFR1L regulates mitochondrial morphology.

Tilokani L, Russell FM, Hamilton S, Virga DM, Segawa M, Paupe V, Gruszczyk AV, Protasoni M, Tabara LC, Johnson M, Anand H, Murphy MP, Hardie DG, Polleux F, Prudent J. Sci Adv. 2022 Nov 11;8(45):eabo7956. doi: 10.1126/sciadv.abo7956. Epub 2022 Nov 11. PMID: 36367943

6. Phosphorylation and activation of cAMP-dependent protein kinase by phosphoinositide-dependent protein kinase.

Cheng X, Ma Y, Moore M, Hemmings BA, Taylor SS. Proc Natl Acad Sci U S A. 1998 Aug 18;95(17):9849-54. doi: 10.1073/pnas.95.17.9849. PMID: 9707564

7. The Importance of the Right Framework: Mitogen-Activated Protein Kinase Pathway and the Scaffolding Protein PTPIP51.

Dietel E, Brobeil A, Gattenlöhner S, Wimmer M. Int J Mol Sci. 2018 Oct 22;19(10):3282. doi: 10.3390/ijms19103282. PMID: 30360441

8. A cold-stress-inducible PERK/OGT axis controls TOM70-assisted mitochondrial protein import and cristae formation.

Latorre-Muro P, O'Malley KE, Bennett CF, Perry EA, Balsa E, Tavares CDJ, Jedrychowski M, Gygi SP, Puigserver P. Cell Metab. 2021 Mar 2;33(3):598-614.e7. doi: 10.1016/j.cmet.2021.01.013. Epub 2021 Feb 15. PMID: 33592173

9. APN-mediated phosphorylation of BCKDK promotes hepatocellular carcinoma metastasis and proliferation via the ERK signaling pathway

Mengying Zhai, Zixia Yang, Chenrui Zhang, Jinping Li, Jing Jia, Lingyi Zhou, Rong Lu, Zhi Yao & Zheng Fu

Cell Death & Disease volume 11, Article number: 396 (2020)

10. Role of the Pyruvate Dehydrogenase Complex in Metabolic Remodeling: Differential Pyruvate Dehydrogenase Complex Functions in Metabolism.

Park S, Jeon JH, Min BK, Ha CM, Thoudam T, Park BY, Lee IK. Diabetes Metab J. 2018 Aug;42(4):270-281. doi: 10.4093/dmj.2018.0101. PMID: 30136450
